# Microalgal protein AstaP is a potent carotenoid solubilizer and delivery module with a broad carotenoid binding repertoire

**DOI:** 10.1101/2021.08.05.455261

**Authors:** Yury B. Slonimskiy, Nikita A. Egorkin, Thomas Friedrich, Eugene G. Maksimov, Nikolai N. Sluchanko

## Abstract

Carotenoids are lipophilic substances with many biological functions, from coloration to photoprotection. Being potent antioxidants, carotenoids have multiple biomedical applications, including the treatment of neurodegenerative disorders and retina degeneration. Nevertheless, the delivery of carotenoids is substantially limited by their poor solubility in the aqueous phase. Natural water-soluble carotenoproteins can facilitate this task, necessitating studies on their ability to uptake and deliver carotenoids. One such promising carotenoprotein, AstaP (Astaxanthin-binding protein), was recently identified in eukaryotic microalgae, but its structure and functional properties remained largely uncharacterized. By using a correctly folded recombinant protein, here we show that AstaP is an efficient carotenoid solubilizer that can stably bind not only astaxanthin but also zeaxanthin, canthaxanthin, and, to a lesser extent, β-carotene, i.e. carotenoids especially valuable to human health. AstaP accepts carotenoids provided as acetone solutions or embedded in membranes, forming carotenoid-protein complexes with an apparent stoichiometry of 1:1. We successfully produced AstaP holoproteins in specific carotenoid-producing strains of *Escherichia coli*, proving it is amenable to cost-efficient biotechnology processes. Regardless of the carotenoid type, AstaP remains monomeric in both apo- and holoforms, while its rather minimalistic mass (∼20 kDa) makes it an especially attractive antioxidant delivery module. *In vitro*, AstaP transfers different carotenoids to the liposomes and to unrelated proteins from cyanobacteria, which can modulate their photoactivity and/or oligomerization. These findings expand the toolkit of the characterized carotenoid-binding proteins and outline the perspective of the use of AstaP as a unique monomeric antioxidant nanocarrier with an extensive carotenoid-binding repertoire.

## Introduction

Carotenoids are highly lipophilic substances performing various biological functions across all kingdoms of life [1-3]. Peculiar chemical properties allow carotenoids to efficiently absorb light and play critical roles in light harvesting, photoprotection and coloration, as well as in preventing and counteracting oxidative stress. With rare exceptions, being chemically modified carotenoid derivatives such as ethers or carbohydrated forms, most carotenoids are poorly soluble in aqueous systems, which makes their biomedical applications quite limited without the development of new approaches. One of such approaches is suggested by nature, as many organisms including cyanobacteria, invertebrates, and microalgae have evolved multiple unrelated water-soluble carotenoid-binding proteins that efficiently solubilize carotenoids and adapt them to the aqueous phase. The most well-studied examples include the coloration-related crustacyanins from crustacean shells [4] as well as the photoprotective two-domain Orange Carotenoid Protein (OCP) [5, 6] and natural homologs of its N- and C-terminal protein domains, i.e. Helical Carotenoid Proteins (HCPs) [7] and C-terminal domain homologs (CTDHs) [8] from cyanobacteria.

β-Crustacyanin, a dimeric block in the α-crustacyanin oligomer, was one of the first carotenoid-binding proteins successfully studied by crystallography [4]. According to the crystal structure (PDB ID 1GKA), this protein dimer binds two molecules of astaxanthin (AXT) [4], which originates from the diet of crustaceans and can be a source of carotenoids for the following members of the trophic chain [9-11]. AXT binding to α-crustacyanin is associated with a significant color change, or the so-called bathochromic shift, from red-orange for free AXT (∼470 nm) to blue-purple in the protein complex (∼632 nm), which underlies the coloration mechanism of these animals [4, 12]. OCP is a unique photosensory protein using a single noncovalently bound ketocarotenoid molecule as a cofactor [5], which endows OCP with photoactivity [13]. Upon high light illumination, OCP reversibly undergoes a structural transformation from the dark-adapted orange OCP^O^ form with a compact protein structure, in which the two protein domains share the embedded ketocarotenoid molecule, to the light-adapted red OCP^R^ form with the carotenoid migrated deep into the N-terminal domain (NTD) and the NTD and CTD domains spatially separated [14-16]. Such dramatic structural reorganization enables OCP^R^ interaction with the cyanobacterial antenna complexes, phycobilisomes, to perform the so-called non-photochemical quenching and dissipating the excess absorbed energy into heat, thereby preventing photodestruction of the photosynthetic apparatus [6, 17-20]. Apart from this phycobilisome quenching activity, OCP is a potent singlet oxygen quencher [21].

As mentioned above, besides the full-size OCP, many cyanobacteria contain genes encoding individual proteins homologous to the NTD and CTD of OCP [7, 8, 17, 22]. Moreover, while some cyanobacteria have OCP and the homologs of its domains simultaneously, others may lack either the homologs or the full-size OCP [22, 23]. This significantly complicates understanding of the roles of those OCP homologs in particular. It is known that HCPs have very similar all-α structures [7, 24], can quench phycobilisome fluorescence and/or reactive oxygen species (ROS) [25], and play a role of carotenoid carriers [26, 27]. The C-terminal domain homologs (CTDHs) of OCP can quench only ROS [8]. Thanks to the outstanding ability of CTDHs to extract carotenoids from the membranes, they were proposed to deliver carotenoids to OCP and/or to HCP [8, 28]. In contrast to predominantly monomeric HCPs, upon carotenoid binding, CTDHs adopt homodimeric structures sharing the carotenoid molecule, which is accompanied by a significant bathochromic spectral shift [8, 26-29]. Such spectral changes, reflecting changes in the carotenoid microenvironment, rather unique in every carotenoprotein, has facilitated the discovery in 2017 of the protein-to-protein carotenoid transfer, which could be readily monitored by optical spectroscopy techniques [28, 30]. For instance, canthaxanthin migration from the liposome membranes, where its absorbance maximum is close to 470 nm, to CTDH results in a ∼90-100 nm bathochromic shift (absorbance maximum at 560-570 nm), which is accompanied by color change from orange-yellow to violet-purple [31].

Carotenoid transfer was found to be a multidirectional process. CTDHs can extract carotenoids from the membranes and redistribute carotenoids to HCP or OCP, but can also extract carotenoids from HCP and even from OCP upon photoactivation of the latter [27]. This finding has allowed us to discover the ability of CTDH to efficiently deliver echinenone to the liposomes and the membranes of mammalian cells [31]. Moreover, such carotenoid transfer by CTDH could cause a significant antioxidative effect by preventing ROS accumulation [31]. These findings proved that the natural water-soluble carotenoproteins can be robust antioxidant nanocarriers and delivery modules with various practical applications [31].

Of note, such a success in studying OCP and related proteins became possible thanks to the ability to obtain the corresponding holoproteins upon expression in carotenoid-producing *Escherichia coli* cells. The first publications in 2015 [32] and 2016 [33], describing such *E. coli* strains, initiated an avalanche-like accumulation of articles reporting structural and mechanistic aspects of those proteins [8, 16, 18, 26-31, 33, 34].

For other carotenoproteins apart from β-crustacyanin and OCP, there is relatively little structural information available. While their carotenoid-binding functions have been well-documented, structural-functional studies were limited by the necessity to isolate protein-carotenoid complexes from natural sources. Among such examples are the Carotenoid-binding protein from silkworm [35], the Echinenone-binding protein from sea urchin [36], Pectenovarin from scallop ovary [37], and AXT-binding protein Ovorubin from apple snail [38, 39].

Recently, Kawasaki and co-workers have discovered a novel water-soluble AXT-binding protein, which they called AstaP, by isolating it from unicellular eukaryotic algae [40]. Although initially these microorganisms were identified as *Scenedesmus* Ki-4, later it was defined as *Coelastrella astaxanthina* Ki-4 [41], where AstaP accumulated in response to the photooxidative stress and presumably provided microalgae with the exceptional resistance to photodamage by performing ROS-quenching and sunscreen activity [40]. AstaP was described as a secretory protein containing an N-terminal signal peptide for transmembrane secretion, a so-called fasciclin-like homology domain, and several glycosylation sites, presumably required to increase stability and solubility of the protein [40]. Overexpressed in stressed algal cells, AstaP was successfully isolated, N-terminally sequenced and identified to be a representative of a new class of carotenoid-binding proteins homologous to the fasciclin family proteins. Usually, these proteins have roles in cell adhesion, cell proliferation, tumor development, and plant reproduction but lack documented carotenoid-binding ability [40]. Interestingly, despite the presence in microalgae of at least several carotenoids including also adonixanthin, canthaxanthin and lutein, the isolated AstaP complex contained predominantly AXT [40, 42]. Although this suggested high specificity of AstaP toward AXT, the ability of AstaP to form stable complexes with other carotenoids remained unaddressed.

Later on, Kawasaki et al. revealed that AstaP homologs are widely present in several strains of algae and even in some bacterial genomes [43]. They described at least two clades of such proteins differing by their pI values (either acidic or alkalic), the presence or absence of potential N-glycosylation sites, and their typical absorbance spectra, which rationalized the naming of those subgroups as orange and pink AstaPs [44]. While these novel water-soluble carotenoproteins are of great interest for basic and applied science, many of their structural and functional properties remain largely unexplored.

To fill these gaps, in the present study, we aimed at obtaining and characterizing the recombinant AstaP from *C. astaxanthina* Ki-4 in *E. coli*, corresponding to the mature protein lacking the signal peptide and posttranslational modifications. We show that the recombinant protein is well-folded and capable of AXT binding. The use of recombinant apoprotein has allowed us to saturate it with the carotenoid and estimate binding stoichiometry and the absorbance properties of the pure holoform. We show that AstaP forms a stable ∼20 kDa monomer that can bind one molecule of AXT, and that the resultant protein-pigment complex is not photosensory. By using different *E. coli* strains capable of producing specific carotenoids, we show that AstaP can form stable complexes with CAN and ZEA, while complexation with β-carotene (βCar) is much less efficient. Carotenoid binding does not change the oligomeric state of AstaP. By using analytical spectrochromatography, we demonstrate that, while AstaP can deliver all tested carotenoid types to the liposomes, carotenoid transfer to unrelated cyanobacterial proteins is more specific and depends on the nature of carotenoids. We show successful delivery of CAN to the CTDH protein from *Anabaena*, which triggers its oligomerization, and either CAN or AXT to OCP from *Synechocystis*, which renders this protein photoactive. These findings significantly expand the toolkit of characterized water-soluble carotenoproteins and pave the way to their rationalized use in modular antioxidant delivery systems.

## Results

### Obtaining and characterization of recombinant AstaP protein

Most native microalgal AstaP proteins possess on the N terminus a hydrophobic signal peptide for transmembrane secretion (Fig. 1A) [44]. Our initial attempts to obtain recombinant full-length AstaP with the signal peptide in *E. coli* were unsuccessful. Therefore, we designed and produced the construct lacking this N-terminal sequence. To the best of our knowledge, any experimental structural data on AstaP are missing. AstaP contains a central sequence most closely homologous to the so-called fasciclin-like domain (residues 29-166, Pfam 02469), the structure of which has been determined by NMR (PDB ID 1W7D) [45] (Fig. 1A). However, AstaP from *C. astaxanthina* Ki-4 and the fasciclin-like protein are characterized by only 36% identity on a range of 140 aligned residues (including 8% gaps). According to the PONDR prediction [46], AstaP features an intrinsically disordered (ID) N terminus, preceding the fasciclin-like domain, and a small ID prone portion in the C terminus of the protein (Fig. 1B). The middle part of the protein is classified as ordered/folded according to the PONDR prediction [46] as well as according to charge-hydropathy plots (with the absolute mean net charge <0.1 at the mean scaled hydropathy of >0.5, based on calculation using the PONDR webserver (http://www.pondr.com)).

**Fig. 1.**
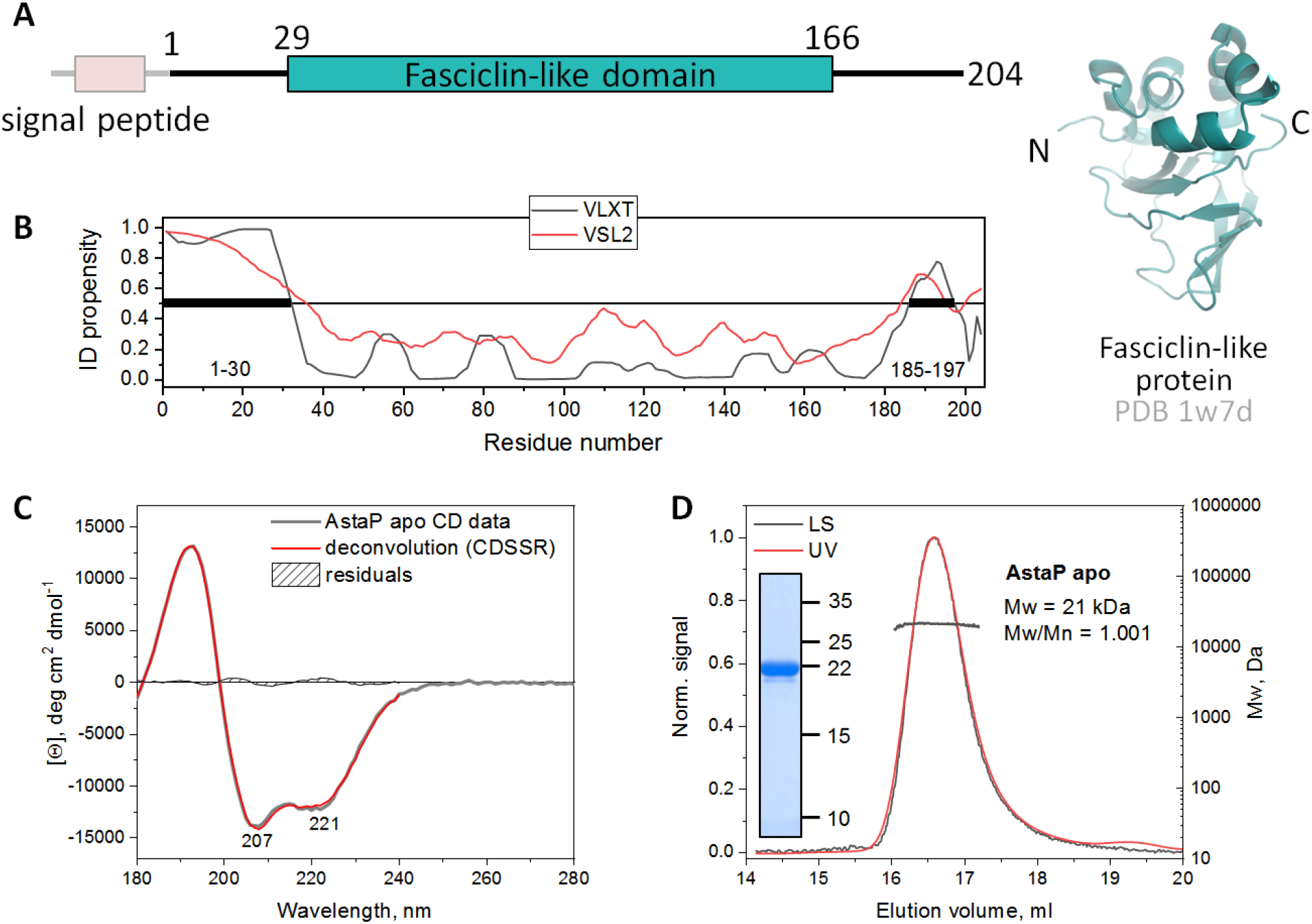
Characterization of the recombinant AstaP apoprotein. A. Schematic representation of the primary structure of AstaP highlighting the fragment used in the present study (1-204). Three-dimensional structure of the fasciclin-like domain, homologous to the central AstaP region (residues 29-166), is shown. B. Per-residue intrinsic disorder propensity of AstaP, predicted by two different PONDR algorithms [46], indicating that the N-terminal portion of the protein is significantly disordered. C. Analysis of the secondary structure of recombinant AstaP using far-UV CD spectroscopy. The experimental spectrum of AstaP was deconvoluted using the DichroWeb server [75], with secondary structure contents indicated in Table 1. The reconstituted CD spectrum and residuals are shown. D. SEC-MALS analysis of the oligomeric state of the recombinant AstaP using a Superdex 200 Increase 10/300 column at a 0.8 ml/min flow rate. Average Mw values across the peak and polydispersity index (Mw/Mn) are indicated, alongside the SDS-PAGE analysis (Mw markers are indicated in kDa).

The far-UV CD spectrum of AstaP has two characteristic minima at 207 and 221 nm (Fig. 1C). Spectral deconvolution suggests the presence of a mixture of secondary structure elements (Table 1). α-Helical content of AstaP (207 residues), assessed by two independent approaches (32-38%) matches that (∼32%) predicted by structure calculation using the most powerful recently published algorithms, AlphaFold2 [47] and RoseTTAFold [48] (see Table 1). These numbers are somewhat higher than the α-helical content in the 1W7D structure of the fasciclin-like protein (24%), suggesting the presence of extra α-helical structures in AstaP beyond its fasciclin-like domain. The contribution from β-strands assessed via deconvolution of the CD spectrum (13%) is less closely but still reasonably predicted by structure modeling (∼20%). The CD data ascribe the most significant contribution to unstructured regions (50%). This is in good agreement with that in the 1W7D structure of the fasciclin-like protein (55%) or structural models built by AlphaFold2 or RoseTTAFold (∼47-48%) (Table 1).

**Table 1.** Secondary structure analysis for the recombinant AstaP apoform based on the far-UV CD spectroscopy.

| Composition | Greenfeld-Fasman [79] | DichroWeb * | AlphaFold2 model ** | RoseTTAFold model ** | 1W7D structure (fasciclin-like protein) *** |
| --- | --- | --- | --- | --- | --- |
| α-helices | 32-38% | 36% | 66/207 (31.9%) | 67/207 (32.3%) | 33/137 (24%) |
| β-strands | - | 14% | 42/207 (20.3%) | 42/207 (20.3%) | 29/137 (21%) |
| unstructured | - | 50% | 99/207 (47.8%) | 98/207 (47.3%) | 75/137 (55%) |

\*The best fit (NRMSD 0.022) is obtained using the CDSSTR algorithm of DichroWeb [75].
\*\*Structural models of AstaP were calculated by either AlphaFold2 [47] or RoseTTAFold [48] using the multiple sequence alignments option. The secondary structure content of the resulting structural models was analyzed by STRIDE [80].
\*\*\* Analyzed by STRIDE [80].

Such a predominance of unstructured regions apparently contradicts the modest ID propensity of AstaP predicted from its sequence (Fig. 1A). Interestingly, the largely unstructured (∼55%) fasciclin-like protein (PDB 1W7D) is still considered well-ordered by PONDR [46]. For example, the absolute mean net charge of the fasciclin-like protein is 0.1 at the mean scaled hydropathy of >0.5. These observations put AstaP in a cohort of proteins whose amino acid content makes it a difficult target for ID prediction algorithms, requiring accurate structural studies.

According to SEC-MALS, recombinant AstaP forms monodisperse particles with Mw of 21 kDa, which closely matches the calculated Mw for protein monomer (21.6 kDa) and its electrophoretic mobility (∼22 kDa) on SDS-PAGE (Fig. 1D). It is worth noting that native AstaP (calculated Mw of 21.2 kDa) eluted on SEC as a 42-44 kDa protein and had an electrophoretic mobility of around 33 kDa, with the discrepancy ascribed to multisite N-glycosylation and oligomeric state proposed to be a monomer [40]. Besides the more accurate Mw determination and confident oligomeric state assignment for the recombinant protein, we note that AstaP is sufficiently stable even in the absence of N-glycosylation-related modifications, which are not installed in *E. coli*.

### Astaxanthin-binding properties of recombinant AstaP

AstaP was first described as the AXT-binding protein whose accumulation in microalgae cells is triggered by desiccation, salt stress and high light [40]. Therefore, we first asked if the recombinant protein was capable of AXT binding (Fig. 2A). Stock solution of commercial all-trans AXT in acetone (0.8 mM) was added to 9 uM AstaP apoprotein to provide for increasing amounts of AXT. AXT concentration in acetone was determined using the molar extinction coefficient of 130,491 M^-1^ cm^-1^ at 477 nm calculated based on data from [49]. Preliminary experiments showed that the addition of acetone up to the final concentration of as high as 12% did not cause AstaP apoprotein aggregation and precipitation, as no protein loss was detected on subsequent SEC profiles. Under these conditions, we incubated AstaP/AXT mixtures for 30 min at room temperature to allow for the complex formation. After removal of unbound AXT by centrifugation, the supernatant was analyzed by spectrochromatography. Elution profiles followed by 460 nm absorbance revealed the single peak corresponding to AstaP(AXT) complex with an apparent Mw of ∼18 kDa, whose amplitude increased with increasing AXT added to AstaP (Fig. 2B). Unbound AXT was utterly insoluble in protein solution and did not appear in any SEC fraction after centrifugation, which supported the conclusion that the stable AstaP(AXT) complex was reconstituted *in vitro*. This notion was further supported by a ∼6 nm bathochromic shift relative to that of AXT in acetone of the AXT absorbance spectrum in the AstaP peak recorded in the course of elution (Fig. 2C). The maximum at 483 nm precisely matches that for the native protein [40]. Using the diode array detector, we were able to monitor spectral changes of the AstaP(AXT) complex upon titration (Fig. 2D). This allowed us to build a pseudo-binding curve that saturated at the Vis/UV ratio of ∼3 (Fig. 2E), reflecting saturation of the apparent AXT-binding capacity of AstaP. To assess binding stoichiometry, protein concentration in the SEC fraction of the reconstituted saturated AstaP(AXT) complex was determined by the Bradford assay, while AXT concentration was determined by acetone extraction following lyophilization. This analysis revealed that within the complex, the apparent molar extinction coefficient of AXT is 4.9 times higher than that of the protein (22,460 M^-1^ cm^-1^ at 280 nm), accounting to ∼110,000 M^-1^ cm^-1^ (at 483 nm). This number is in the same order of magnitude as that of AXT in petroleum ether (143,000 M^-1^ cm^-1^ at 470 nm), acetone (130,491 M^-1^ cm^-1^ at 477 nm), or benzene (118,097 M^-1^ cm^-1^ at 489 nm) (as calculated based on data from [49]). Carotenoid extraction and determination of the molar extinction coefficient of AstaP(AXT) at 483 nm supported the binding stoichiometry of 0.98 AXT per 1 AstaP. Noteworthily, the native isolated AstaP featured a Vis/UV absorbance ratio of only 1.8 [40], which in comparison with our value observed at saturation *in vitro* (∼3), might indicate a mixture of apo and holoproteins. Carotenoid extraction in the original study [40] enabled the determination of the carotenoid-protein molar ratio of 0.75. Therefore, the consensus carotenoid binding stoichiometry of AstaP is close to 1:1. Given AXT insolubility in aqueous solutions, the apparent efficiency of AstaP binding to AXT, and our ability to prepare quite concentrated solutions of the AstaP(AXT) complex, AstaP appears a very efficient carotenoid solubilizer.

**Fig. 2.**
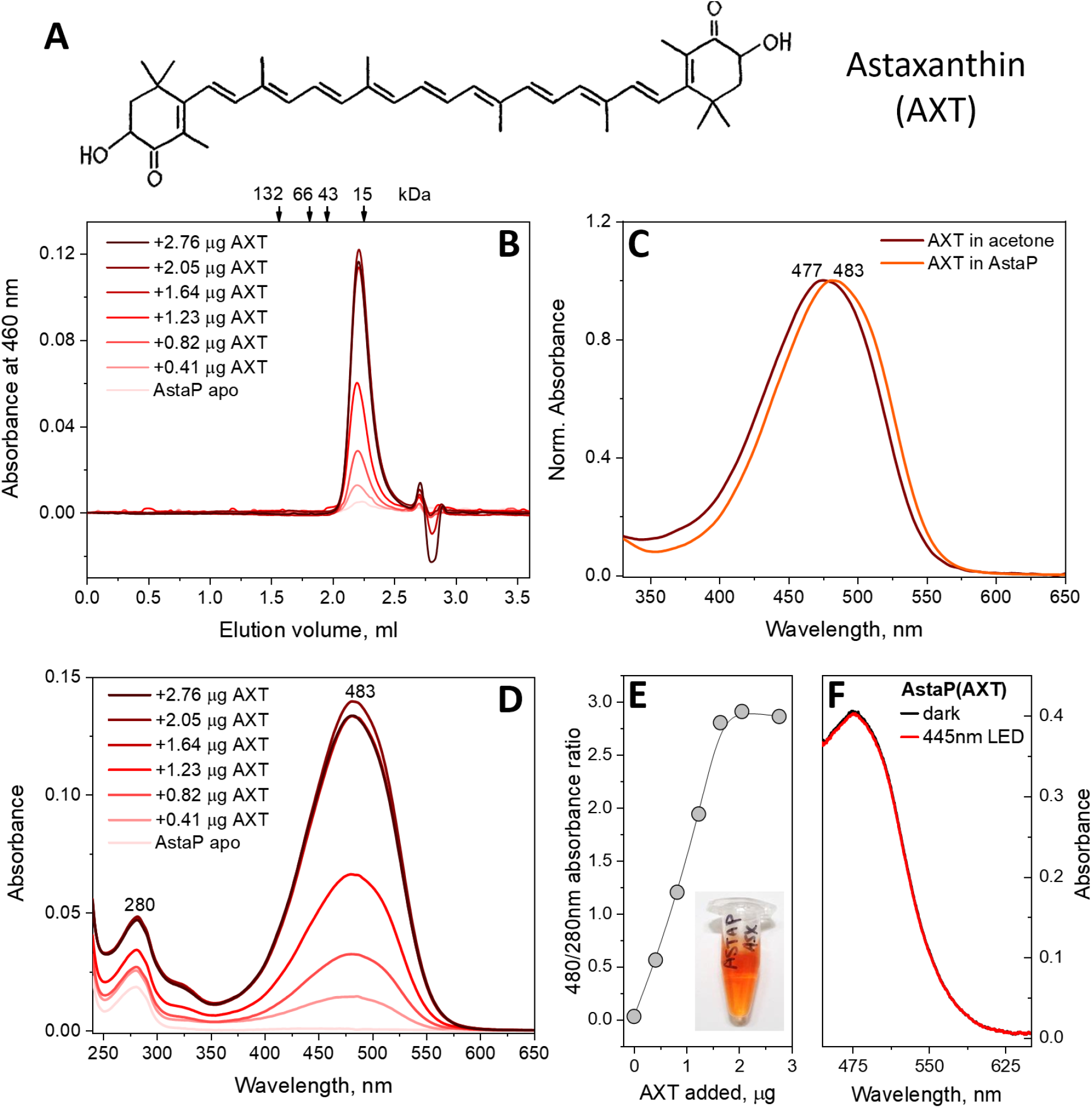
Recombinant AstaP efficiently binds astaxanthin (AXT) from acetone solution, yielding water-soluble monomeric carotenoprotein. A. Structural formula of AXT. B. Size-exclusion spectrochromatography profiles of the reconstituted AstaP(AXT) complex obtained after pre-incubation of recombinant AstaP with various amounts of AXT (indicated in micrograms) added as acetone solution. SEC traces at 460 nm are shown, while all spectra in the range of 240-650 nm were recorded using the diode array detector (DAD). C. Absorbance spectra of AXT in acetone and in protein. D. Absorbance spectra of the AstaP(AXT) complex formed upon the titration experiment, as retrieved from the SEC peak maximum using the DAD data. E. Saturation of AstaP by AXT built as an increase of the Vis/UV absorbance ratio as a function of increasing amounts of AXT added. F. The absorbance spectrum of the saturated AstaP(AXT) complex before and after blue LED illumination.

AstaP is known to be upregulated in response to high light stress [40, 44]. Since it is the first time that the stable AstaP holoprotein is at hand at high concentrations and excellent purity, we tested whether the protein would show any signs of photoactivity, similar to the Orange Carotenoid Protein from cyanobacteria [13, 50]. However, even intense light exposure (10 min at 200 mW, blue LED) of the AstaP(AXT) sample did not result in any changes in the absorbance spectrum (Fig. 2F). This disfavors the hypothesis that AstaP holoprotein is a photosensory protein, unless its photoactivated intermediates relax at a very high rate.

### AstaP efficiently extracts different xanthophylls from biological membranes

We then asked if not only AstaP can take different carotenoids *in vitro* but also extract them from biological membranes. To test this possibility, we mixed the purified apoprotein with suspensions of *E. coli* membranes obtained from strains genetically modified to produce either ZEA, CAN, or βCar (Fig. 3A) (See Materials and Methods for more details). These mixtures were incubated for 30 min at room temperature and then overnight at 4 °C, centrifuged, and then soluble material was analyzed by spectrochromatography. This resulted in the appearance on the chromatograms of strong visible absorbance for ZEA or CAN co-eluting with the AstaP peak, for the first time indicating efficient formation of AstaP complexes with these carotenoids (Fig. 3B). On the contrary, almost no visible absorbance was detected for βCar, which indicated rather limited βCar binding or extraction efficiency by AstaP.

**Fig. 3.**
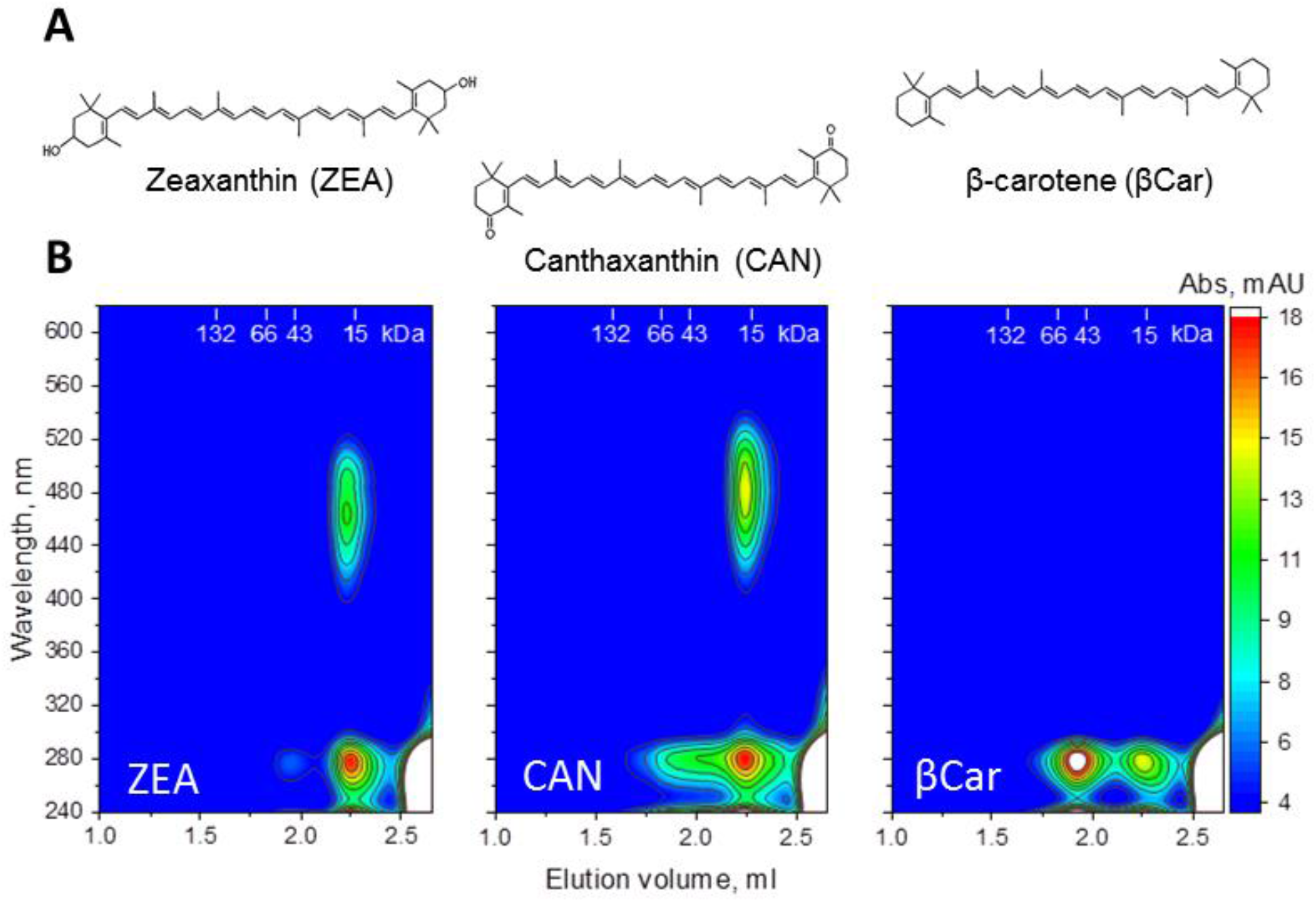
Recombinant AstaP is capable of carotenoid extraction from *E. coli* membranes. A. Structural formulae of the carotenoids individually expressed in specific strains of *E. coli* and provided to AstaP as membrane suspensions. B. Spectrochromatograms of AstaP-carotenoid complexes showing the accumulation of ZEA (left panel) or CAN (middle panel) in the protein fraction (apparent Mw of ∼18 kDa, positions of protein markers (kDa) are indicated on the top). Under the same conditions, β-carotene (right panel) is not bound by AstaP. Amplitudes of absorbance are color-coded according to the scale on the right.

Having found that AstaP does not need other factors to extract carotenoids from membranes, we tried to obtain holoproteins by directly expressing AstaP in the carotenoid-producing strains of *E. coli*. As above with the apoprotein, the construct was supplied with the N-terminal His_6_-tag cleavable by specific 3C protease to facilitate protein purification (Fig. 4A). A combination of chromatography steps enabled fast and efficient protein purification (Fig. 4B), aided by carotenoid absorption in the visible spectral region (Fig. 4C). Purification from ZEA- or CAN-producing cells has allowed us to obtain preparative amounts of AstaP holoproteins (Fig. 4D), whose carotenoid contents were confirmed by acetone extraction followed by thin-layer chromatography (Fig. 4E). Such analysis for the AstaP(ZEA) complex revealed that both *E. coli* membranes and the protein preparation contain almost exclusively ZEA. In the case of AstaP(CAN), a remarkable accumulation of CAN takes place – while *E. coli* membranes contained CAN and a large fraction of βCar, no βCar could be extracted from AstaP, which contained CAN exclusively (Fig. 4E). Although this clearly indicated that AstaP prefers CAN over βCar, we tested if βCar can still be bound by AstaP upon large-scale expression in *E. coli* cells producing only βCar. This indeed yielded a colored protein fraction whose absorbance spectrum had a characteristic vibronic structure and a ∼10 nm bathochromic shift relative to βCar absorbance in acetone (Fig. 5A). While thin-layer chromatography confirmed the presence of βCar in this preparation (Fig. 5B), the Vis/UV absorbance ratio of <0.3 suggested limited efficiency of βCar binding.

**Fig. 4.**
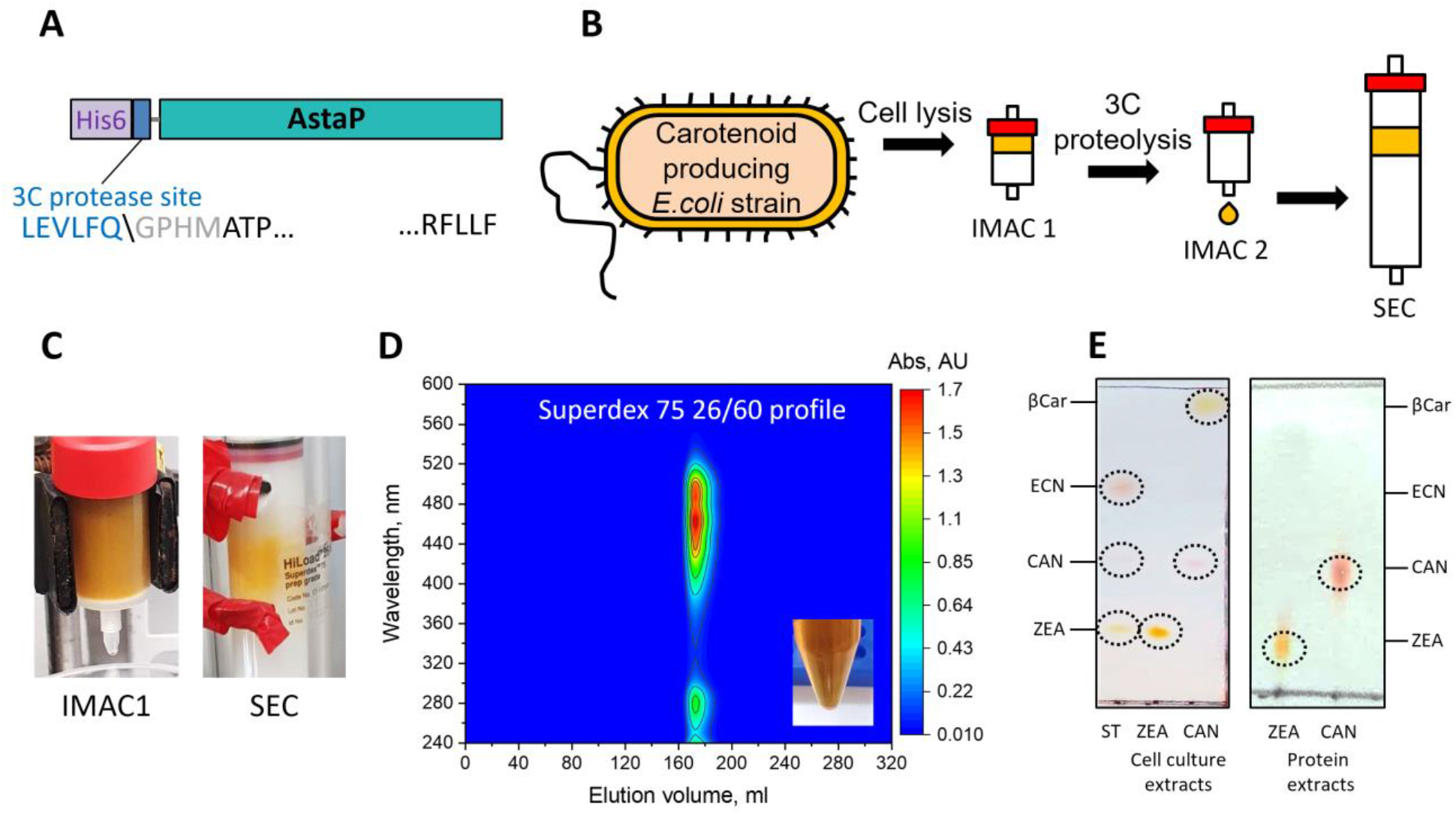
Obtaining AstaP-carotenoid complexes in *E. coli* cells producing specific carotenoids. A. Schematic showing the design of AstaP facilitating its purification via a cleavable His_6_-tag. B. A scheme showing the process of purification of AstaP-carotenoid complexes using subtractive immobilized metal-affinity chromatography (IMAC1 and IMAC2) and size-exclusion chromatography (SEC). C. Appearance and color of the columns during the IMAC and SEC steps for AstaP(ZEA). D. A typical spectrochromatogram of the AstaP(ZEA) complex obtained using a Superdex 75 26/600 column and DAD data. The appearance and color of the sample are shown in the inset. E. Thin-layer chromatography of the carotenoid content in *E. coli* cells at the end of the cultivation (left) and in the purified AstaP(ZEA) and AstaP(CAN) complexes (right).

**Fig. 5.**
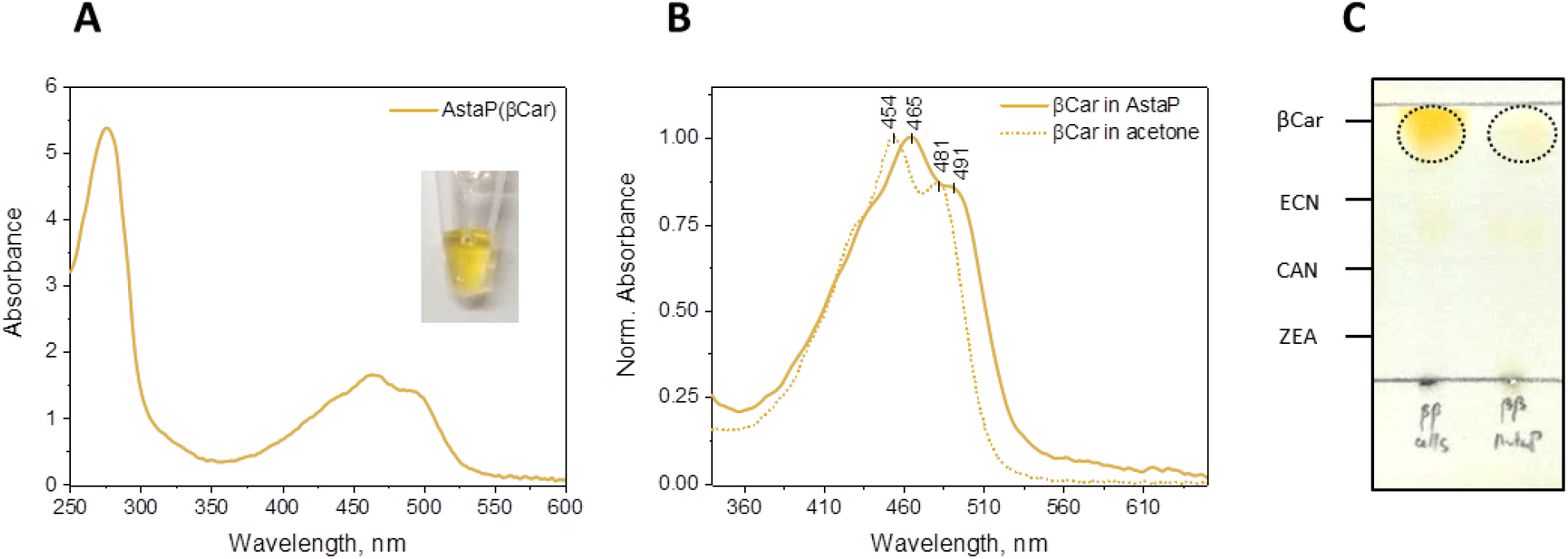
Recombinant AstaP is capable of β-carotene (βCar) binding, although with limited efficiency. A. Absorbance spectrum of the AstaP(βCar) complex. The sample color is shown in the inset. B. Absorbance spectra of βCar in acetone and in the AstaP(βCar) complex. C. Thin-layer chromatogram showing βCar presence in the AstaP fraction and in the *E. coli* cells at the end of cultivation. Characteristic positions of different carotenoids (β-carotene, echinenone, canthaxanthin, zeaxanthin) are indicated on the left.

The marked difference in binding efficiency of βCar and xanthophylls (i.e, oxygenated carotenoids AXT, ZEA, and CAN) can indicate that the hydroxyl- or keto-groups of those xanthophylls make contacts with the protein which stabilize the complex or at least take part in the initial recognition of these ligands by AstaP.

The AstaP(CAN) and AstaP(ZEA) complexes were further characterized by spectrochromatography coupled to MALS (Fig. 6). Single peaks were observed on the elution profiles, and the Mw distribution unequivocally indicated monodisperse sets of particles for either AstaP(CAN) or AstaP(ZEA). However, to determine the absolute Mw values for these species, the use of the weight extinction coefficient calculated for the apoprotein (ε^0.1%^_280nm_=1.04) was incorrect due to the significant carotenoid absorbance in the UV range. To take this into account, we determined the concentration of AstaP(CAN) or AstaP(ZEA) complexes using the Bradford assay, which is insensitive to the presence of carotenoids. This has allowed us to correct the extinction coefficients at 280 nm for the corresponding holoproteins to 1.44 and 1.75, respectively, and to more accurately determine Mw as 21 [AstaP(CAN)] and 22 kDa [AstaP(ZEA)]. Both numbers are in excellent agreement with the calculated masses for the corresponding AstaP holoprotein monomers (∼22 kDa), which indicates that carotenoid binding with AstaP does not interfere with its oligomeric state.

**Fig. 6.**
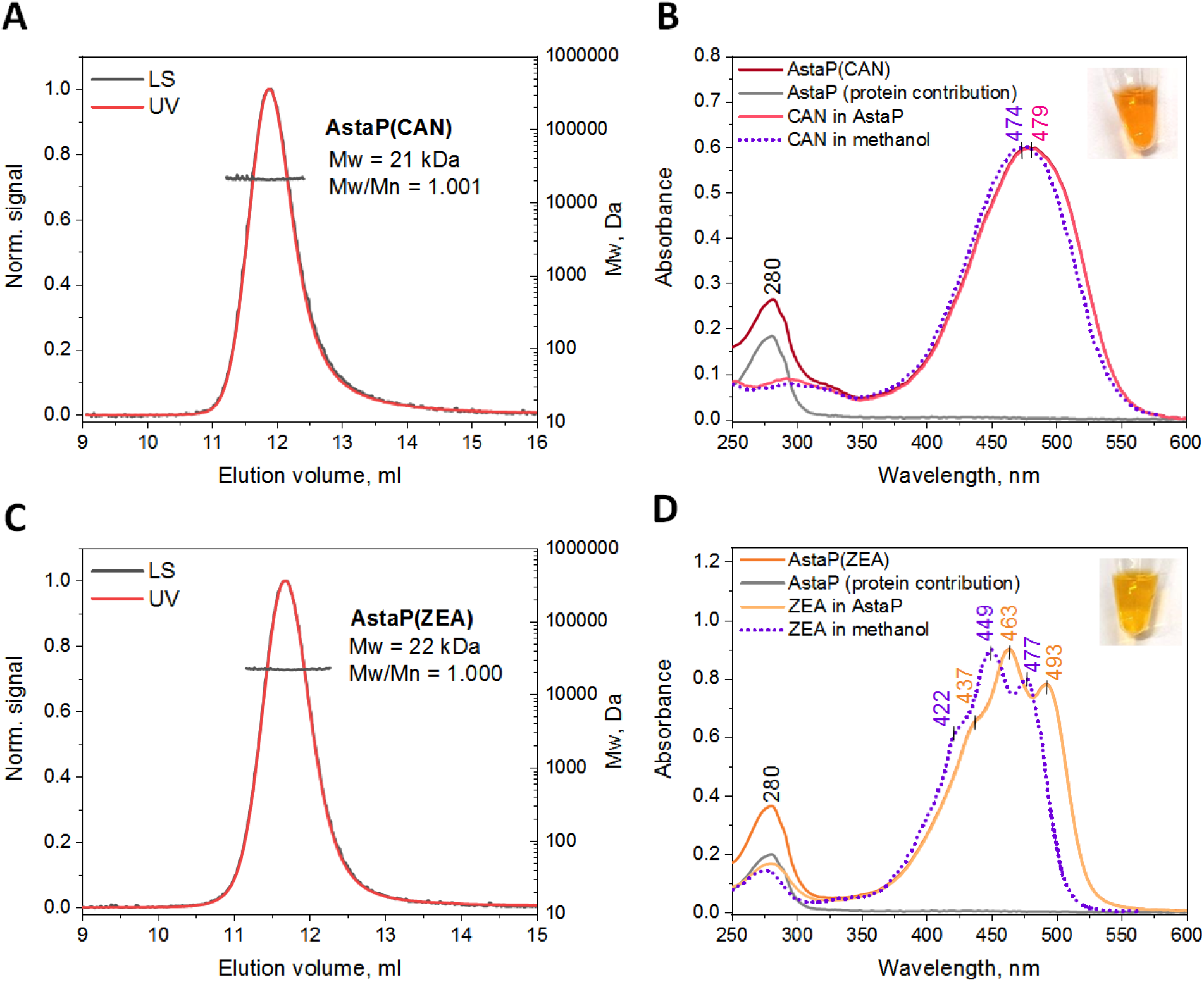
Properties of the AstaP(CAN) and AstaP(ZEA) complexes obtained from *E. coli* cells producing the corresponding carotenoids. A and C show SEC-MALS profiles for the AstaP(CAN) and AstaP(ZEA) complexes using a Superdex 75 10/300 column at a 0.8 ml/min flow rate. C and D show absorbance spectra of CAN (or ZEA) in methanol and in complex with AstaP. The latter is shown along with the decomposition into protein and carotenoid spectra. Sample colors are seen in the insets. Decomposition was done using the corrected extinction coefficients for the holoforms calculated based on the independent protein concentration measurement by the Bradford assay [74].

Using diode array detection, we retrieved absorbance spectra from the maxima of the SEC peaks, which revealed bathochromic shifts upon binding to AstaP for both CAN and ZEA, compared to their absorbance in methanol (Fig. 6C and D). CAN and ZEA spectra in methanol were retrieved from www.lipidbank.jp [51]. The shift upon AstaP binding for ZEA (∼15 nm) was much more significant than for CAN (∼5 nm), which may indicate different conformations of hydroxylated and ketolated carotenoids in the AstaP complex.

Independent protein concentration determination using the Bradford assay has allowed us to estimate the contribution of carotenoids in the UV absorbance and reconstruct full absorbance spectra of ZEA and CAN within their complexes with AstaP. Of note, the decomposed contributions of ZEA and CAN absorbance in the UV region within the AstaP complex very closely match the absorbance peaks for these carotenoids in methanol (Fig. 6B and D).

Similar to AstaP(AXT), neither AstaP(ZEA) nor AstaP(CAN) could be photoactivated by even intense blue LED illumination (data not shown), further supporting the notion that AstaP is not itself a photosensory protein.

### AstaP delivers different xanthophylls to liposomes via transient interaction

We next questioned whether AstaP can deliver its carotenoids to membrane fraction and tested this hypothesis using model liposomes. Previous work has demonstrated the usefulness of analytical spectrochromatography (ASESC) for monitoring protein-protein and protein-liposome carotenoid transfer [27, 30, 31, 52].

In a similar setup, we incubated the purified AstaP holoproteins embedding different xanthophylls (ZEA or CAN or AXT) with liposomes and analyzed the results of carotenoid transfer by ASESC with full-spectrum absorbance detection of the eluate (Fig. 7). The conditions used have allowed us to baseline separate fractions of large liposomes and small proteins and thereby assess spectral characteristics of either independently of each other.

**Fig. 7.**
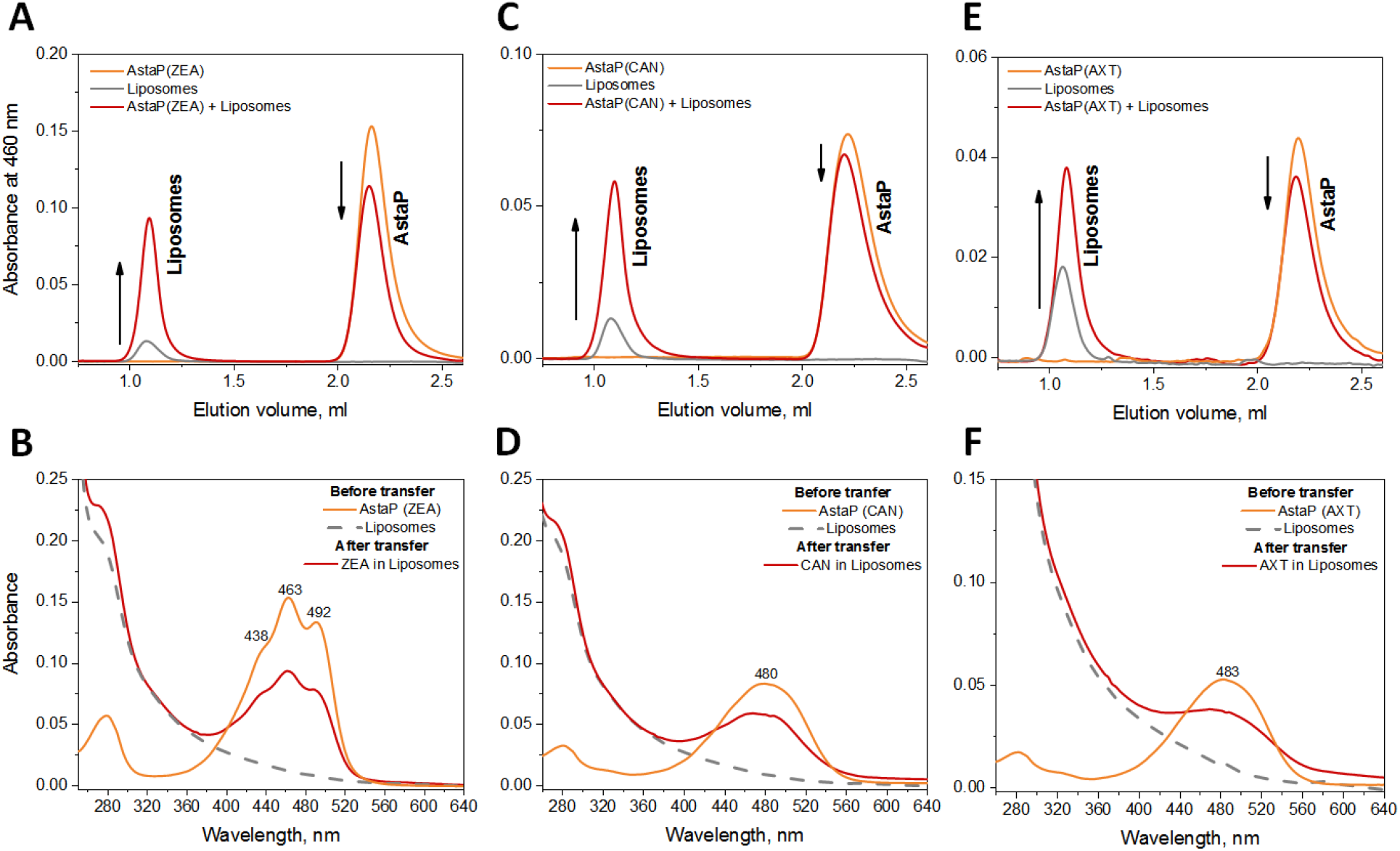
AstaP delivers xanthophylls into liposomes. A, C, E. SEC profiles followed by absorbance at 460 nm from a Superdex 200 Increase 5/150 column at a 0.45 ml/min flow rate showing the positions of liposomes and AstaP holoforms before and after carotenoid transfer. Arrows indicate changes associated with carotenoid transfer from AstaP to liposomes. B, D, F. Absorbance spectra at the AstaP holoform fraction and at the liposomal fraction retrieved from DAD data and showing carotenoid migration to the liposomes, with the associated spectral changes. Note high Rayleigh scattering at the liposome fraction.

In all cases, we observed partial loss of carotenoids from the monomeric AstaP fraction and a concomitant appearance of the characteristic carotenoid absorbance in the liposome fraction (Fig. 7A, C, E). For all three carotenoids, their spectral shapes (e.g, the presence or absence of vibronic structure) were largely preserved upon the translocation from protein to liposomes (Fig. 7B, D, F). In agreement with our SEC-MALS data, carotenoid transfer did not affect the monomeric status of AstaP. This situation is different from that known for the *Anabaena* CTDH protein, which undergoes the monomer-dimer transition upon carotenoid binding [8, 26, 27].

We hypothesized that AstaP can have an affinity to the liposome membranes and tested this hypothesis by fractionation of the AstaP-liposome mixtures according to sizes, with the subsequent SDS-PAGE analysis (Fig. 8A and B). This experiment showed that, although most of the AstaP holoprotein mixed with liposomes remained in the protein fraction, a small amount was detected in the liposome fraction, indicating transient interaction. Unfortunately, all our attempts to analyze co-elution of liposomes with the AstaP apoprotein by SEC failed due to the ability of the latter preparation to provoke instant liposome precipitation even at 150 mM NaCl.

**Fig. 8.**
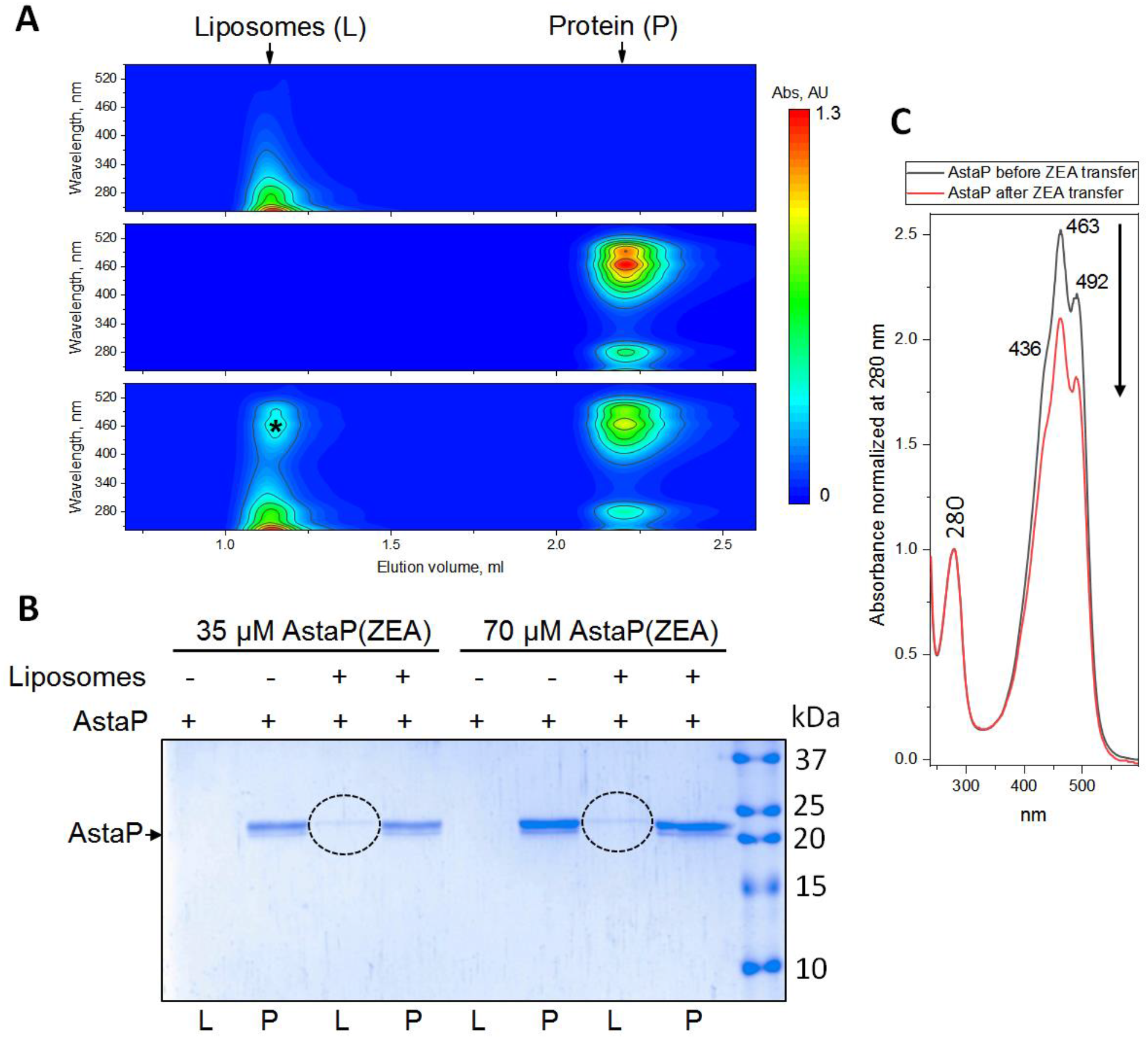
Transient interaction of AstaP with liposomes. A. Spectrochromatograms of liposomes (top), 70 μM AstaP(ZEA) (middle), or their pre-incubated mixture (bottom) obtained on a Superdex Increase 5/150 column using DAD. Liposome (L) and protein (P) fractions are marked by arrows. Asterisk indicates the appearance of ZEA absorbance in the liposome fraction after carotenoid transfer from AstaP(ZEA). Color scale to the right corresponds to absorbance levels, shown with isolines on the plots. B. SDS-PAGE analysis of the protein redistribution between L and P fractions at two different AstaP concentrations used in the experiment (as indicated), such as shown on panel A for 70 μM AstaP. Dashed ovals mark the appearance of the minor quantity of AstaP in the liposome fraction. AstaP position on the gel is shown by the arrow. C. Absorbance spectra of AstaP(ZEA) before and after carotenoid transfer to liposomes measured at the position of the protein peak, normalized to absorbance at 280 nm. Note that the Vis/UV absorbance ratio changes (indicated by arrow) upon carotenoid translocation to the liposomes.

Very importantly, despite the co-elution of the AstaP protein with the liposomes, the appearance of the visible absorbance in this fraction could not be fully accounted by adhesion of the AstaP holoprotein on the membranes and meant *bona fide* carotenoid translocation from protein into the membrane phase. Indeed, if normalized at 280 nm, the absorbance spectra of AstaP(ZEA) before and after mixing with the liposomes underwent the decrease of the Vis/UV ratio associated with the carotenoid loss from the protein fraction (Fig. 8C).

### AstaP transfers ketolated xanthophylls to cyanobacterial Orange Carotenoid Protein

Having a portfolio of AstaP holoproteins with different ligands at hand, we were interested to study the ability of AstaP to deliver cartenoids to other carotenoproteins.

First, we wanted to test if AstaP is able to give carotenoids to the apoform of the Orange Carotenoid Protein, which is the photoactive 35-kDa protein responsible for the photoprotection mechanism in cyanobacteria [6, 17, 20]. The photoactivity of OCP, i.e., its ability to reversibly undergo the OCP^O^ → OCP^R^ transition under actinic light, depends on the nature of the bound carotenoid molecule [53], which could be instrumental for the assessment of the carotenoid delivery from AstaP. It is known that a molecule of ketocarotenoid such as echinenone or 3-hydroxyechinenone or canthaxanthin is required for OCP photoactivity, whereas ZEA binding does not render OCP photoactive [53]. A recent study has detected OCP photoactivity in the presence of a mixture of CAN and AXT [54]; however, the experimental settings used left the question of OCP photoactivity with exclusively AXT unresolved.

We mixed the apoform of *Synechocystis* OCP with various AstaP-carotenoid complexes and analyzed the outcome of the potential protein-protein carotenoid transfer by ASESC. Under our experimental settings, we achieved an excellent spatial separation of the reactants and products of carotenoid transfer to analyze absorbance spectra of the corresponding fractions individually. Apparent Mw values for the peaks were estimated from column calibration.

In the case of CAN transfer, we observed the appearance of the peak with the intermediate position (∼34 kDa) between those of the OCP apoprotein (∼62 kDa) and AstaP(CAN) (∼18 kDa), and a concomitant redistribution of carotenoid between AstaP and this new peak (Fig. 9A, peak 2). Only a minor peak with absorbance in the visible region appeared at the position of the OCP apoprotein (Fig. 9A, peak 1). Previous studies have shown that, like OCP^R^, the OCP apoprotein has an expanded tertiary structure with the two protein domains detached, and in this state is prone to homodimerization, whereas carotenoid binding within the interdomain tunnel causes protein compaction via domain re-association, thereby decreasing its effective hydrodynamic size [14, 30, 33, 55]. In perfect agreement with these notions, we observed that CAN migration from AstaP (∼18 kDa) causes compaction of the expanded OCP apoprotein (∼62 kDa) into ∼34-kDa species with the pronounced vibronic structure characteristic of the compact orange OCP (Fig. 9A and B, peak 2). Along with this major product of CAN transfer from AstaP to OCP, we have detected a minor product characterized by absorbance maximum of 503 nm and a hydrodynamic size similar to that of the OCP apoprotein (Fig. 9A and B, peak 1). This species most likely represents the expanded OCP-carotenoid complex with the detached protein domains with the N-terminal domain binding CAN, while its spectrum with the absorbance maximum of 503 nm likely reflects poor separation of this peak from that of compact orange OCP(CAN) and hence partial cross-contamination.

**Fig. 9.**
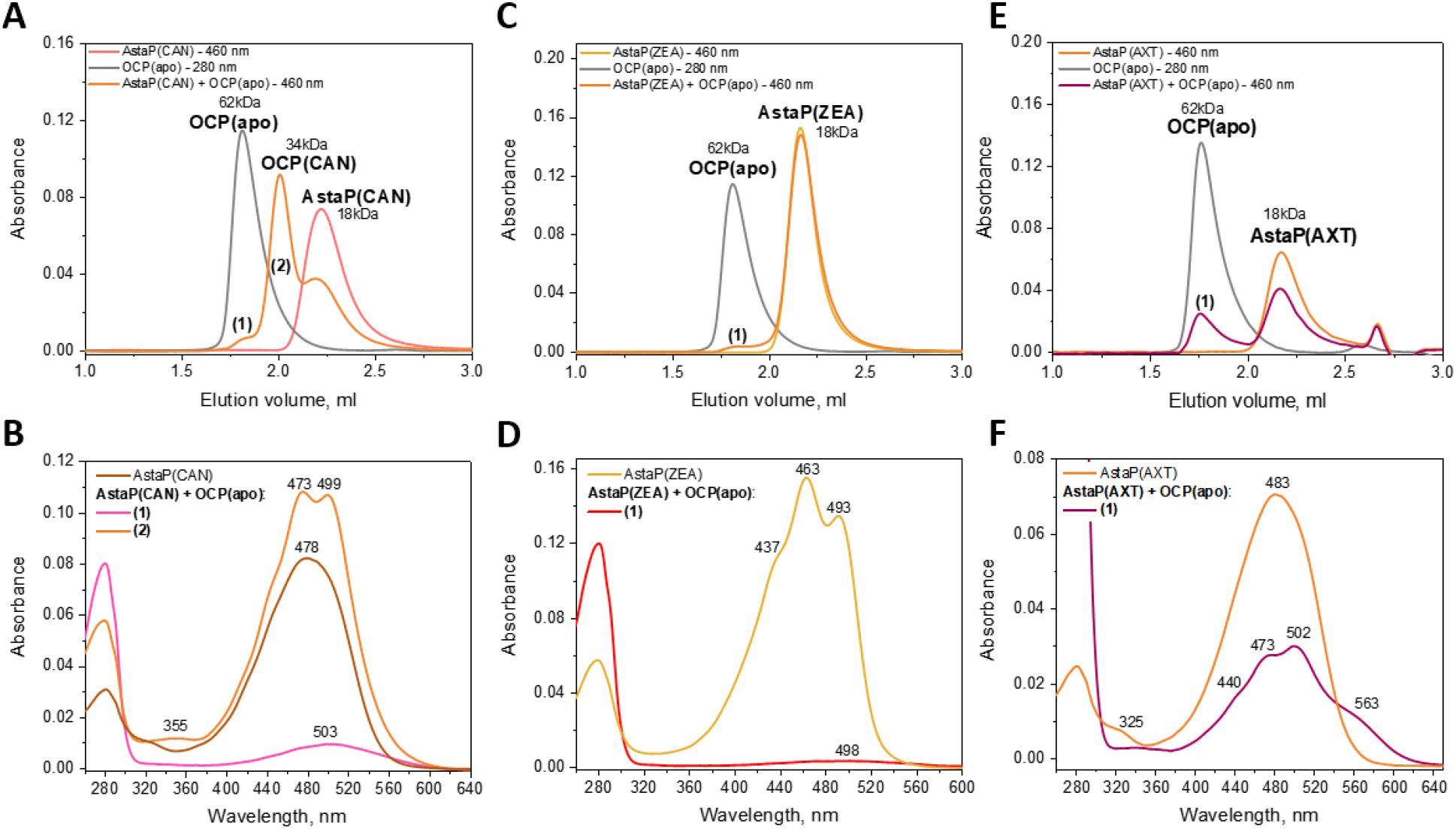
Xanthophyll transfer from AstaP to the OCP apoprotein studied by analytical spectrochromatography. Individual AstaP holoproteins containing either CAN, ZEA, or AXT, or individual OCP apoprotein, or the corresponding AstaP/OCP mixtures were pre-incubated for at least 30 min at room temperature and then loaded on a Superdex 200 Increase 5/150 column at a 0.45 ml/min flow rate upon monitoring full absorbance spectrum of the eluate. A, C, E represent SEC profiles for the results of CAN (A), ZEA (C), or AXT (E) transfer, where the main SEC peaks are assigned and labeled. Newly appeared peaks with absorbance in the visible region are marked either (1) or (2) for each case, and the absorbance spectra corresponding to their maxima are shown in panels B, D, F for CAN, ZEA, or AXT containing species, respectively. The absorbance spectrum of the corresponding AstaP holoform (i.e., carotenoid donor in the experiment) is shown for comparison.

Under similar conditions, we observed almost no transfer of ZEA from AstaP to the OCP apoprotein (Fig. 9C), with only a very small fraction of the newly formed species with the expanded hydrodynamic size and absorbance maximum of around 500 nm (Fig. 9C and D, peak 1). This finding was unexpected because it was previously shown that OCP can bind ZEA in specific cyanobacterial strains lacking ketocarotenoids, which, however, rendered protein non-photoactive [53]. Our result may indicate that AstaP has a much higher affinity to ZEA than OCP and does not release this carotenoid.

We could not exclude that the 3,3’-hydroxyl groups of ZEA, absent in CAN, could disfavor ZEA binding to OCP, which would explain the sharp contrast compared to transfer of ketocarotenoid CAN from AstaP to OCP. Therefore, we also studied the possibility of AXT transfer, which possesses both 4,4’-ketogroups and 3,3’-hydroxyl groups simultaneously (Fig. 2A). Intriguingly, we observed substantial AXT transfer from AstaP to OCP leading to the formation of the OCP species with the vibronic structure (three peaks at 440, 473, and 502 nm) and a shoulder at around 563 nm, likely representing a mixture of protein forms (Fig. 9E and F, peak 1). To the best of our knowledge, this is the first report on spectral characteristics of the OCP species containing purely AXT. We speculate that the shoulder at 563 nm likely reflects the formation of the OCP(AXT) dimers which share AXT between their CTDs, similar to what was earlier reported for OCP(CAN) [28, 34]. The appearance of the vibronic structure in the OCP(AXT) peak can imply the similarity to a better described OCP(CAN) and hence, the photoactivity of this species.

### Photoactivity of the OCP species formed by carotenoid transfer from AstaP

The findings that AstaP can transfer different carotenoids to OCP prompted us to test if the OCP species formed are photoactive, i.e. can respond to blue LED illumination (actinic light) by changes of the absorbance spectrum [13, 50].

The mixture of AstaP(CAN) and OCP apoprotein was allowed to equilibrate until carotenoid transfer ended and then was illuminated by 445 nm LED to induce photoactivation. As expected, such treatment led to profound spectral changes with the increased absorbance at around 550 nm and loss of the vibronic structure (Fig. 10A). As described elsewhere, such spectral changes can be used to monitor the OCP photocycle, which is a temperature-sensitive process [13, 50]. We confirmed that the OCP species formed after CAN transfer from AstaP displays reversible photoactivation and temperature-dependent relaxation in the dark, being much longer at 10 °C than at 30 °C (Fig. 10B).

**Fig. 10.**
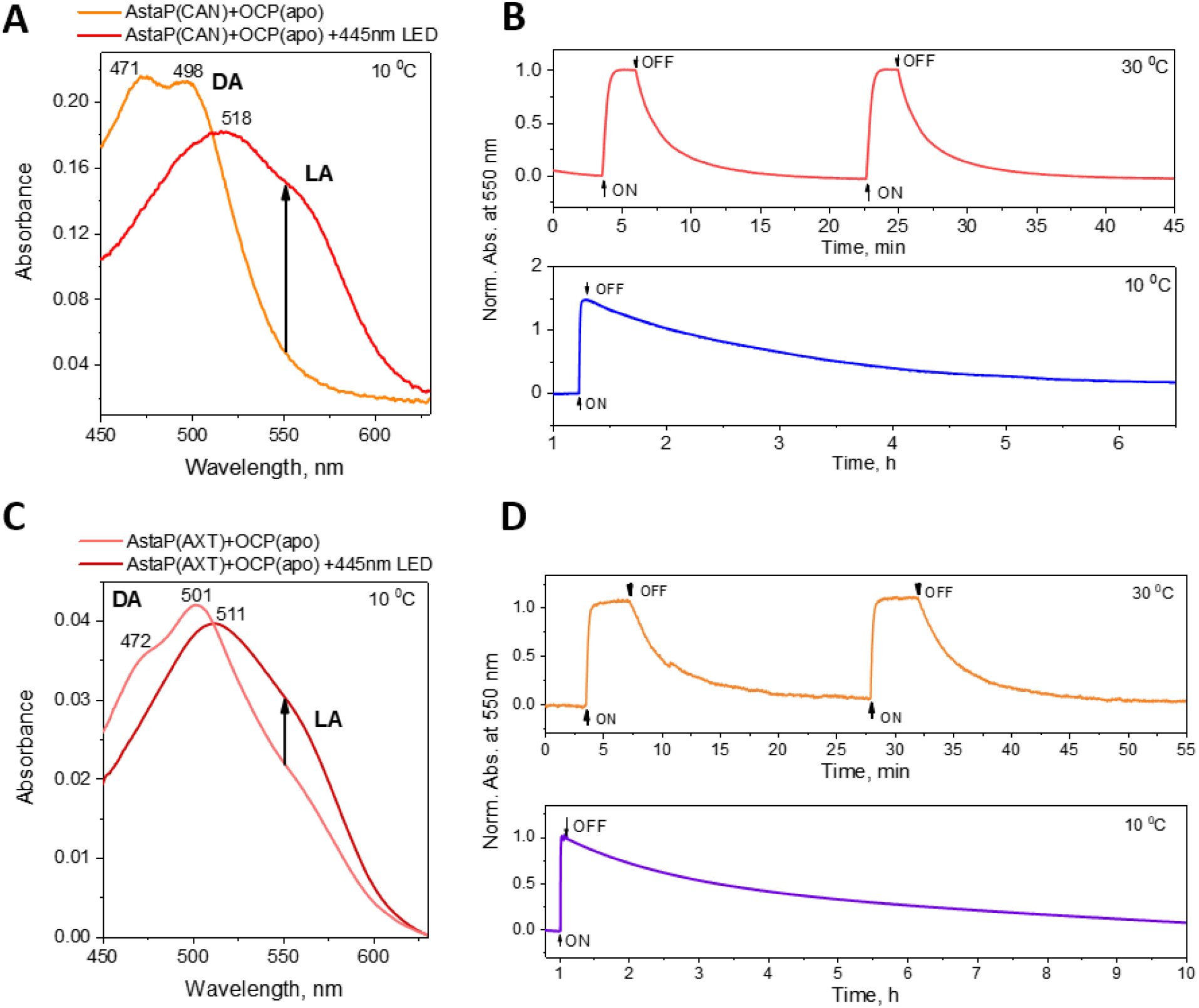
Photoactivity of OCP after carotenoid transfer from AstaP. Canthaxanthin (A) or astaxanthin (C) transfer from AstaP to the OCP apoprotein yielded photoactive OCP species whose transition from dark-adapted (DA) to light-adapted (LA) forms was triggered by 445 nm LED (indicated by arrows) at the indicated temperature. Since the AstaP(AXT) sample added to OCP contained acetone, the OCP(AXT) species were first isolated by SEC prior to the photoactivity test. B. Normalized changes of absorbance at 550 nm were used to monitor at either 30 °C or 10 °C the photocycle of the OCP(CAN) species formed after carotenoid transfer from AstaP. Note the difference in the time scales (min and h for 30 and 10 °C, respectively). D. Normalized changes of absorbance at 550 nm were used to monitor at 30 °C or 10 °C the photocycle of the OCP(AXT) species formed after carotenoid transfer from AstaP and separation on SEC. Time points designating switching actinic light on and off are indicated on panels B and D.

Intriguingly, we could also observe that the much less characterized OCP(AXT) species obtained after AXT transfer from AstaP, is also photoactive (Fig. 10C). Moreover, we could reconstitute the photoactive OCP(AXT) complex simply by adding AXT as the acetone solution and the following buffer exchange (data not shown). In both scenarios, the vibronic structure of the absorbance spectrum was observed for the dark-adapted protein, while blue LED illumination caused spectral transformation into a form lacking any vibronic structure and having increased far red-shifted absorbance (Fig. 10C). A shoulder at ∼563 nm, present in the dark-adapted absorbance spectrum of OCP(AXT), seemed irresponsive to the photoactivation, in favor of the hypothesis that it corresponds to the dimeric OCP species sharing AXT via the CTD-CTD interaction. Just like OCP(CAN), OCP(AXT) species displayed the reversible photocycle, with the back-relaxation even more significantly dependent on temperature – at 10 °C it took 1.5-2 times longer for OCP(AXT) than for OCP(CAN) (Fig. 10D). In line with the previously proposed theory [56-58], this observation can indicate that for AXT the expanded OCP structure requires more attempts to re-establish the dark-adapted compact form, probably due to the presence of the 3,3’-hydroxyl groups in AXT, which can interfere with acquiring proper orientation of this bulkier carotenoid.

### AstaP transfers specific xanthophylls to cyanobacterial CTDH protein, triggering its oligomerization

We have previously characterized a powerful carotenoid delivery protein module derived from *Anabaena variabilis* CTDH [31]. This protein efficiently transferred carotenoids to the membranes of liposomes and mammalian cells, thereby counteracting oxidative stress in the latter [31]. However, the practical use of this protein may be limited by its carotenoid specificity, as, to our knowledge, CTDH was shown to form stable complexes only with echinenone and canthaxanthin [27, 31]. Moreover, while CTDH was capable of extracting from membranes both echinenone and canthaxanthin, it could efficiently deliver only echinenone [31].

Since broadening of the repertoire of carotenoids accommodated by CTDH would be of great biomedical use, and given that AstaP can handle several carotenoids valuable to human health, we studied the possibility of carotenoid transfer from AstaP holoproteins to the CTDH apoprotein. In the absence of carotenoids, *Anabaena* CTDH exists as a mixture of ∼18 kDa monomers and ∼38 kDa dimers, with the monomer being prevailing (Fig. 11A). This is consistent with the previous reports [27, 31]. Incubation of the CTDH apoprotein with AstaP(CAN) resulted in a gradual color change from orange to violet-purple, suggesting protein-protein carotenoid transfer. On the ASESC profile followed by visible absorbance, we observed a dramatic redistribution of the peaks – the decreasing amplitude of the AstaP(CAN) peak and the appearance of two new peaks (Fig. 11A). Absorbance spectra of these peaks with the maxima at 559 (peak 2) and 564 nm (peak 1) (Fig. 11B) and their positions on the elution profile (apparent Mw of 38 and 57 kDa) suggested the CTDH dimers and higher-order oligomers containing CAN (probably trimers and tetramers stabilized by distinct interfaces observed in the CTDH apoform structure, 6FEJ [26, 27, 59]). The absorbance maximum in peak 1 is far red-shifted (∼564 nm) compared to CAN absorbance in AstaP (∼478 nm). This would agree well with the previously observed ∼570 nm absorbance maximum for the *Tolypothrix* CTDH tetramer [59]. Nevertheless, due to incomplete spatial separation of peaks 1 and 2 (Fig. 11A) and to partial overlap with peak 2 with the larger amplitude and the absorbance maximum of 559 nm, the *bona fide* absorbance maximum of the peak 1 CTDH oligomers (∼57 kDa) can be even farther red-shifted. Of note, this spectrum features a shoulder at around 600 nm. Therefore, CAN transfer from AstaP is associated with a dramatic change of the carotenoid environment, reflected in a 85-100 nm bathochromic shift.

**Fig. 11.**
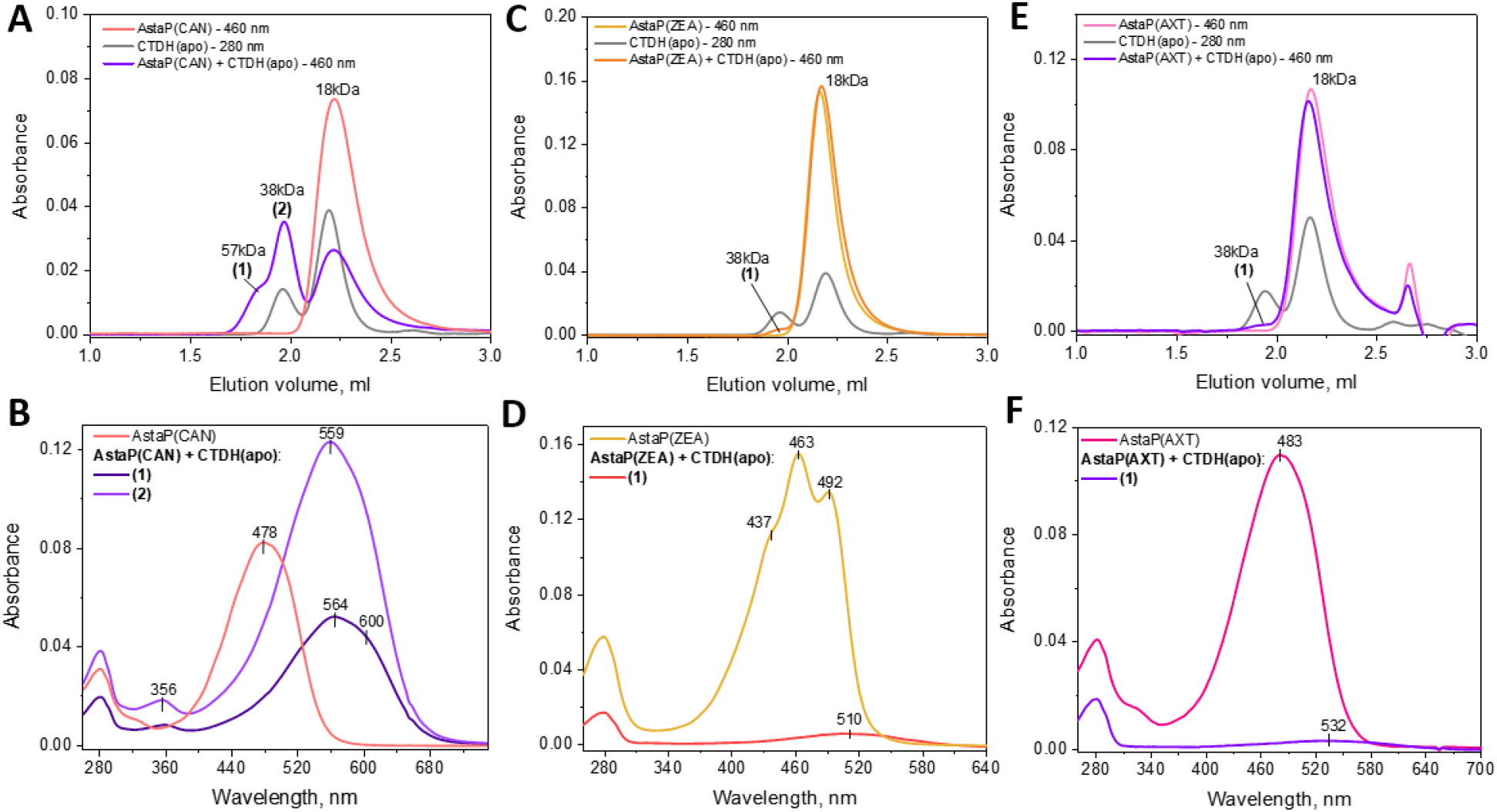
Xanthophyll transfer from AstaP to the CTDH apoprotein studied by analytical spectrochromatography. Individual AstaP holoproteins containing either CAN, ZEA, or AXT, or individual CTDH apoprotein, or the corresponding AstaP/CTDH mixtures were pre-incubated for at least 30 min at room temperature and then loaded on a Superdex 200 Increase 5/150 column at a 0.45 ml/min flow rate upon monitoring full absorbance spectrum of the eluate. A, C, E represent SEC profiles for the results of CAN (A), ZEA (C), or AXT (E) transfer. Newly appeared peaks with absorbance in the visible region are marked either (1) or (2) for each case, and the absorbance spectra corresponding to their maxima are shown in panels B, D, F for CAN, ZEA, or AXT containing species, respectively. The absorbance spectrum of the corresponding AstaP holoform (i.e., carotenoid donor in the experiment), is shown for comparison.

In striking contrast, AstaP transferred almost no ZEA to CTDH as the elution profile contained nearly unchanged peaks corresponding to the CTDH species and AstaP(ZEA) (Fig. 11C). A minor peak with absorbance in the visible region (maximum at ∼510 nm) could be detected at the position of the CTDH dimers. A very similar situation was observed in the case of AXT. Under conditions used, only a minimal peak with the visible absorbance spectrum (maximum at ∼532 nm) was detected at the position of the CTDH dimers (Fig. 11E).

These results suggest that the affinity of AstaP to ZEA or AXT significantly exceeds that of CTDH. At the same time, we were able to detect the formation of principally new holoforms of the dimeric CTDH protein embedding ZEA or AXT. This suggests that by adjusting experimental parameters, one could shift the equilibrium to ensure the desired yield of certain carotenoproteins.

Due to the coincidence of the SEC peaks of the CTDH and AstaP monomers, we cannot rule out that some amount of ZEA (or AXT) was still transferred to the monomeric fraction of CTDH. However, due to the unchanged absorbance spectrum in this part of the elution profile, this probability is rather low.

Therefore, AstaP can accommodate different carotenoids and deliver some of them to acceptor carotenoproteins.

## Discussion

In this work we successfully designed, produced in *E. coli*, and characterized recombinant carotenoprotein AstaP from a microalga *C. astaxanthina* Ki-4, where it is overexpressed in response to stress conditions requiring maximal photoprotection [40, 42]. N-terminal sequencing of the protein isolated from the native source indicated that the hydrophobic signal peptide was lacking [40, 43]. Therefore, our design of the recombinant AstaP protein matched the original protein, except for the extra residues GPHM… present in our construct after cleaving off the His-tag.

Native isolated AstaP, the so-called AstaP-orange1, was extensively N-glycosylated [42]. This protein presumably resides in the outer surface of the plasma membrane of the AstaP-expressing microalgae cells and apparently withstands severe desiccation and salt stress. Under these conditions, protein solubility may become a problem, especially for proteins embedding hydrophobic substances like carotenoids. It is very likely that N-glycosylation reduces this problem by increasing the solubility and stability of AstaP. The representatives of the AstaP-pink clade do not seem to contain glycosylation sites, yet they have acidic pI values, which may somehow compensate for the absence of glycosylation in keeping protein sufficiently stable [44]. Of note, the pink AstaPs are predicted to be targeted to the endoplasmic reticulum and should function under milder conditions [44].

Nevertheless, recombinant AstaP-orange1, studied in this work, proved stable and soluble enough even without glycosylations, which do not occur in *E. coli*. Our CD data and the predictions made here by the latest algorithms based on artificial intelligence and deep learning implemented in AlphaFold2 [47] and RoseTTAFold [48], together suggest that AstaP is a well-folded protein. While these structure prediction algorithms support the previous proposition that AstaP contains a fasciclin-like domain (Pfam 02469), our analysis indicates that protein regions beyond the fasciclin domain may have additional structured elements. In agreement with the CD spectroscopy data, AlphaFold2 [47], RoseTTAFold [48], and PONDR [46] coherently predict the presence of the intrinsically disordered regions (IDRs) especially in the N-terminal portion of the protein. This complicates the structure prediction and also raises the question about the functional role of these IDRs. For instance, AstaP-orange2 does not contain 19 residues of the N-terminal IDR of AstaP-orange1 [44]. Intriguingly, in spite of these IDRs and the presence of up to 50% of unstructured regions as suggested by our CD data, AstaP is still predicted to be an ordered protein by several algorithms of PONDR [46], thereby presenting a curious example of a Janus protein with two faces.

Recombinant AstaP formed monodisperse monomers, which proved fully functional in binding of its cognate ligand, AXT. The maximum (∼483 nm) and shape of the absorbance spectra of native and recombinant proteins closely match. This indicated that native expression is not obligatory for the obtaining of functional AstaP. Moreover, the use of the recombinant apoprotein lacking posttranslational modifications permitted accurate functional analysis. This revealed that AstaP binds AXT with the apparent stoichiometry close to 1:1 at saturation, with the Vis/UV absorbance ratio approaching 3 and being much higher than detected previously for the native protein (1.8) [40]. This lower ratio may probably be explained by partial loss of carotenoids upon isolation or incomplete protein loading by carotenoids *in vivo*. The determination of binding stoichiometry was complicated by the appreciable absorbance of AXT in the UV region, which required careful carotenoid extraction and quantitation. While the AXT binding mechanism to AstaP awaits further elucidation by structural biology techniques, we assume that at least *in vitro* AstaP is saturated upon binding 1 AXT per 1 protein.

In AstaP complex, AXT undergoes a subtle bathochromic shift from 477 nm in acetone to 483 nm in protein. On one hand, this indicates that the protein matrix does affect the conformation of the carotenoid, but, on the other hand, this red shift (∼6 nm) is incomparably smaller than that known for the AXT binding to α-crustacyanin (∼160 nm) [4]. One explanation to this difference may be that AstaP does not significantly affect the effective conjugation length of AXT, possibly because of making only minimal chemical contacts with AXT. AstaP was described by Kawasaki et al as the protein upregulated in response to excessive illumination [40, 42-44], which could indicate that it is a photosensory protein. However, our attempts to detect photoactivation-related changes in the absorbance spectrum of AstaP(AXT) were unsuccessful, suggesting that AstaP upregulation is triggered by different mechanisms rather involving changes in gene expression.

Given that native AstaP was isolated preferentially as the AXT complex even despite several other carotenoids such as lutein, adonixanthin and canthaxanthin were evenly present in microalgal cells [42, 43], the question arose whether AstaP could form stable complexes with other carotenoids as well. In this work, we showed the formation of functional AstaP complexes with AXT, ZEA or CAN (Fig. 12).

**Fig. 12.**
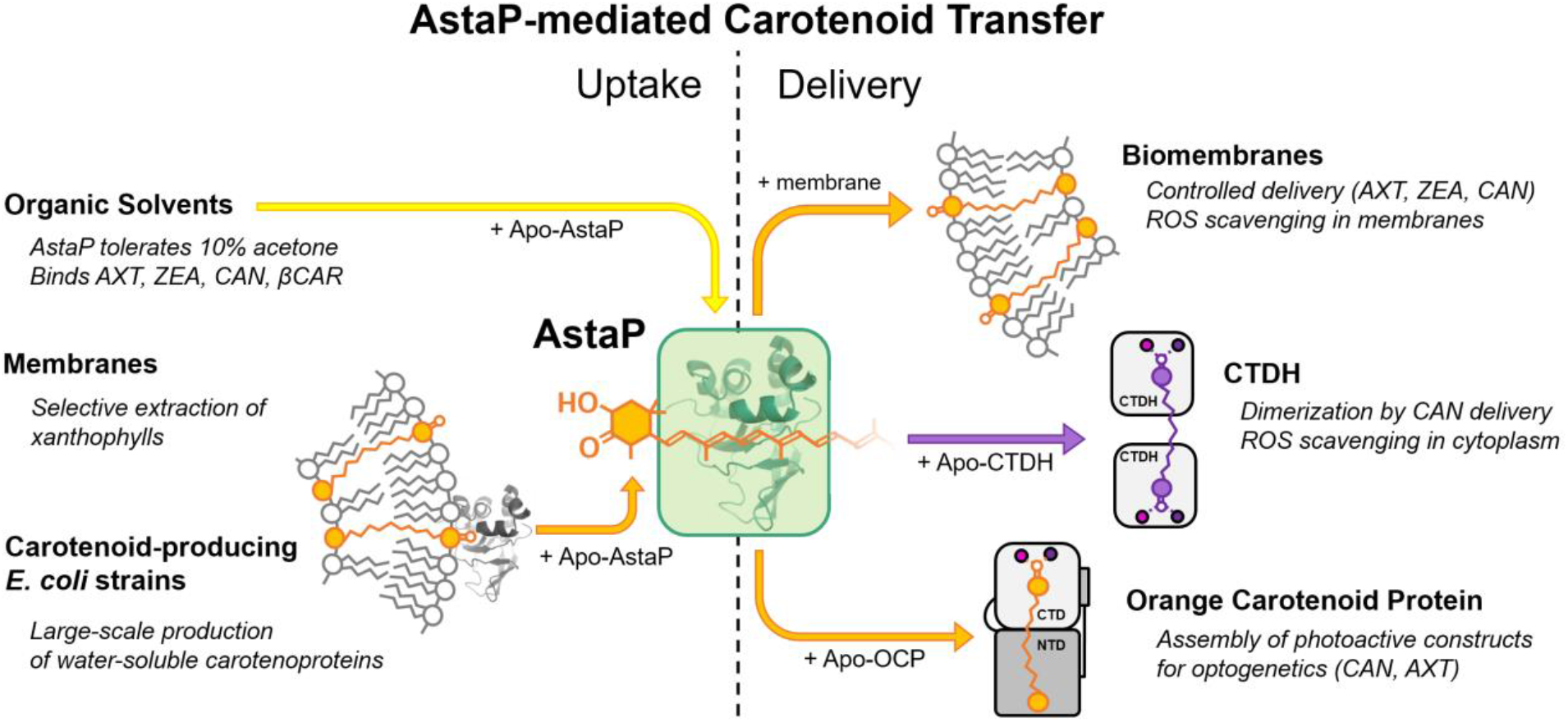
Summary of carotenoid binding and transfer properties of AstaP. See text for details.

Some carotenoproteins, such as most of HCPs from cyanobacteria, require auxiliary proteins supplying carotenoids, whereas others, like cyanobacterial CTDHs and OCPs, can extract carotenoids from membrane compartments on their own [8, 26]. According to our data, recombinant AstaP can very efficiently extract different xanthophylls from biological membranes, not requiring additional protein factors (Fig. 12). While ZEA and CAN could be extracted, yielding colored AstaP holoproteins, βCar extraction proved much less efficient. This may indicate that the presence of the 3,3’-hydroxy- and/or 4,4’-ketogroups on the terminal carotenoid rings may be instrumental for the extraction mechanism by AstaP. Alternatively, the stability of the βCar complex may be weaker than with xanthophylls, or the orientation of βCar in the membranes is unfavorable for the extraction. In any case, recombinant AstaP served as a very efficient carotenoid solubilizer with the capacity to stably accommodate at least several carotenoids relevant to human health (Fig. 12).

Extending our understanding of the AstaP functionality, we demonstrated that it can be readily obtained in *E. coli* strains synthesizing specific carotenoids, whereby AstaP complexes with ZEA or CAN were purified and analyzed (Fig. 12). We used multi-angle light scattering (MALS) coupled to analytical size-exclusion spectrochromatography (ASESC) to simultaneously determine absolute molecular mass for the AstaP-carotenoid complexes and their spectral characteristics. This analysis revealed that carotenoid binding does not affect the preferred monomerization state of AstaP, while the protein caused different bathochromic shifts to the absorbance spectra of ZEA (∼15 nm) and CAN (∼5 nm). The ability of AstaP to stably accommodate ZEA and CAN in addition to its cognate ligand, AXT, may indicate that any of the 3,3’-hydroxy- or 4,4’-ketogroups are sufficient to form productive complexes. If this assumption is correct, one can expect that AstaP can bind even a wider range of xanthophylls.

Next to showing that AstaP is an efficient solubilizer of the biomedically relevant carotenoids AXT, ZEA and CAN, we showed that it can transfer these carotenoids to the liposome membranes via transient interaction (Fig. 12). We suggest that this occurs due to the local electrostatic interactions of the positively charged AstaP (pI = 10.5) with the negatively charged lipid groups even at 150 mM NaCl. These data serve as proof of principle for the successful application of AstaP as an antioxidant nanocarrier. With respect to the broader ligand binding repertoire of AstaP, it can probably be more promising than the previously characterized *Anabaena* CTDH [31]. The use of AstaP can appreciably widen the range of carotenoids to be used for the liposome loading or direct carotenoid delivery to vulnerable cells and tissues (Fig. 12). Recent works discuss a plethora of biological effects and applications for xanthophylls, including ZEA, CAN and AXT [60-71].

Carotenoid binding and release by CTDH are associated with the oligomeric transition, whereas such activity of AstaP leaves its monomeric state unchanged. Given its profound carotenoid binding efficiency and the ∼20 kDa mass, AstaP module is already rather minimalistic and could probably be shortened further at the expense of the variable terminal regions. As mentioned above, AstaP-orange2 from the Oki-4N strain [44] lacks the N-terminal 19 residues PKANATTAKPASTTSTP present within the IDR of AstaP-orange1 studied here. This stretch may be a candidate for removal upon further miniaturization of AstaP.

In addition to the ability to deliver carotenoids to the liposomes, AstaP displayed a remarkable ability to transfer carotenoids to unrelated cyanobacterial proteins OCP and CTDH (Fig. 12). In this study, we confirmed that the OCP(CAN) species are photoactive regardless of the formation mechanism, including CAN delivery from the unrelated AstaP protein. To our knowledge, this is the first report on successful carotenoid transfer between completely unrelated proteins from eukaryotes and prokaryotes. Besides the high efficiency of CAN transfer to both OCP and CTDH, which rendered OCP photoactive and caused oligomerization of CTDH (Fig. 12), we observed AXT transfer to OCP and demonstrated the photoactivity of OCP in the presence of purely AXT for the first time. To the best of our knowledge, this is the first report of AXT transfer between any proteins.

Like for OCP(CAN), we observed a dramatic decrease of the back-conversion of the dark-adapted OCP(AXT) upon decreasing temperature, and in the case of AXT it was even slower than for CAN. This may reflect the necessity of the 3,3’-hydroxygroups to be accommodated within the CTD, requiring additional attempts to form the compact dark-adapted OCP form. Not all carotenoid types accommodated by AstaP could be transferred to recipient proteins. While at least a small fraction of the carotenoid-bound products formed in all cases, the efficiency was limited upon ZEA transfer from AstaP to both OCP and CTDH and upon AXT transfer to CTDH. This may reflect the higher relative affinity of AstaP to these carotenoids or instability of the recipient complexes with these carotenoids.

To sum up, we may conclude that AstaP is a small monomeric carotenoid-binding protein with an unexpectedly broad carotenoid binding repertoire, which can significantly expand the toolkit of the known carotenoproteins for biomedical applications and carotenoid delivery approaches.

## Materials and Methods

### Materials

All-trans-astaxanthin and β-carotene (CAS Numbers: 472-61-7 and 7235-40-7) were purchased from Sigma-Aldrich (USA). Liposomes were prepared according to the Racker method [72] with modifications. For the preparation of liposomes, a buffer solution containing 50 mM KH_2_PO_4_, 2 mM MgSO_4_, pH 7.5 was used. The pH was adjusted to 7.5 with dry potassium hydroxide tablets. Liposomes were prepared from L-α-lecithin isolated from soybeans containing 17% phosphatidylcholine. 40 mg of lecithin was diluted in 1 ml of buffer. The resulting mixture was homogenized in a glass homogenizer until the homogeneous suspension was obtained. The resulting mixture (1 ml) was placed in an Eppendorf tube and sonicated for 40 min on an ultrasonic disintegrator UZDN-2T (Ukrrospribor, Ukraine) at a 22 kHz frequency until the suspension became completely clear. The hydrodynamic size of the liposomes was determined by dynamic light scattering on a ZetaSizer Nano ZS analyzer (Malvern Instruments, UK) to be equal to 80±20 nm. Liposomes were stored at 4 °C and used within two days.

Membranes of specific *E. coli* strain cells with the overexpressed carotenoids βCar, or ZEA, or CAN, used in the present work as carotenoid donors to AstaP, were obtained via washing by lysis buffer of the insoluble membrane fraction of the cultivated *E. coli* cells described below.

Synthesis of βCar was achieved by transformation of *E*.*coli* cells with the pACCAR16ΔcrtX plasmid (chloramphenicol resistance) harboring the contiguous gene cluster consisting of *crtY, crtI, crtB*, and the *crtE* gene from *Erwinia uredovora* for constitutive expression [33]. Synthesis of ZEA was achieved by means of the pACCAR25ΔcrtX plasmid (chloramphenicol resistance) harboring the contiguous gene cluster consisting of *crtY, crtI, crtB, crtZ* and the *crtE* gene from *E. uredovora* for constitutive expression [33].

The *crtE* gene product (a geranylgeranyl-pyrophosphate synthase) forms geranylgeranyl-pyrophosphate from isopentyl-pyrophosphate and farnesyl-pyrophosphate, both of which are naturally formed in *E. coli*. The *crtB* gene product (a phytoene synthase) forms phytoene from two geranylgeranyl-pyrophosphate blocks. The *crtI* gene product (a phytoene desaturase) forms lycopene from phytoene and the *crtY* gene product (a lycopene cyclase) forms βCar from lycopene. The *crtZ* gene product (a βCar hydroxylase) converts βCar in two hydroxylation steps at the 3 and 3’ position of the β rings to ZEA [33].

Synthesis of CAN was achieved by means of combining the pACCAR16ΔcrtX plasmid (chloramphenicol resistance) [33] and the pBAD plasmid (ampicillin resistance) plasmid containing a *crtW* ketolase (adds ketogroups at the 4 and 4’ position of the β rings of βCar) gene from *Anabaena* sp. PCC7210 put under the control of the arabinose inducible promoter araBAD [73], yielding a mixture of CAN (predominant fraction) and βCar (minor fraction). The *crtW* plasmid was a kind gift from Prof. Kai-Hong Zhao (Huazhong Agricultural University, Wuhan, China).

All chemical reagents were of the highest purity and quality available. All aqueous solutions in the study were prepared on the milliQ-quality water (18.2 MΩ/cm).

### Cloning, protein expression, and purification

The apoforms of the full-length wild-type OCP from *Synechocystis* sp. PCC6803 (residues 1-317, Uniprot ID P74102) and the C-terminal domain homolog from *Anabaena variabilis* (residues 1-140, Uniprot ID Q8YMJ3) were produced in *E. coli* BL21(DE3) cells and purified as described earlier [27, 30]. The coding sequence for AstaP from *C. astaxanthina* Ki-4 (residues 1-204, Uniprot ID S6BQ14) was codon-optimized for expression in *E. coli* [58] and synthesized by IDT Technologies (Coralville, Iowa, USA). This sequence was flanked by *Nde*I and *Xho*I sites and was inserted into a pET28-His-3C plasmid (kanamycin resistance) at these sites so that the recombinant protein possessed at the N-terminus the His_6_-tag cleavable by the highly specific human rhinovirus 3C protease. After cleavage the AstaP protein bore extra GPHM… residues on its very N terminus.

The AstaP construct was verified by DNA sequencing (Evrogen, Moscow, Russia) and used to transform chemically competent cells. AstaP apoprotein was expressed using induction by 0.2 mM isopropyl-β-thiogalactoside (IPTG) for 24 h at 30 °C in the presence of kanamycin and purified to electrophoretic homogeneity using a combination of subtractive immobilized metal-affinity (IMAC) and size-exclusion chromatography (SEC). AstaP holoproteins were expressed in the specific *E. coli* cells producing specifically either βCar, or ZEA, or CAN, described above. In the case of CAN-synthesizing strain, crtW expression was induced by the addition of 0.02% L-arabinose. Protein expression was induced by the addition of 0.2 mM IPTG and lasted 24 h to achieve build-up of the desired carotenoids, whose contents in the membrane fraction were tested by thin-layer chromatography following acetone extraction. The purification scheme for the holoproteins was the same as for the apoproteins. Pure protein preparations were stored frozen at −80 °C.

Protein concentrations were determined on a Nanophotometer N80 (Implen, Germany) by absorbance at 280 nm using the sequence-specific extinction coefficients calculated by the Protparam tool in ExPasy. In the case of the holoforms of AstaP, we used the Bradford assay [74] to determine protein concentration independent from the present carotenoids, which was used to correct the corresponding extinction coefficients required for Mw calculation by SEC-MALS. Importantly, in this case, the calibration curve for the Bradford assay was built using the AstaP apoprotein.

### Circular dichroism spectroscopy

AstaP apoprotein (0.7 mg/ml) was dialyzed overnight against 20 mM phosphate buffer, pH 7.2, and then centrifuged for 10 min at 4 °C and 14200 g prior to measurements. Far-UV CD spectra of the sample were recorded at 20 C in the range of 180-260 nm at a rate of 1 nm/min with 0.5 nm steps in 0.01 cm quartz cuvette on a Chirascan circular dichroism spectrometer (Applied Photophysics) equipped with a temperature controller, and then the signal from the buffer filtered through 0.22 μm membrane was subtracted. Secondary structure elements were estimated using the mean residue weight of AstaP equal to 104.4 Da by the DichroWeb server [75] by the CDSSTR algorithm with a set 11 of reference proteins optimized for the range 180-240 nm [76].

### AstaP structure modeling

The tertiary structure of AstaP was calculated by AlphaFold2 [47] and RoseTTAFold [48] algorithms using the custom multiple sequence alignment (MSA) option. MSA was obtained for the query AstaP sequence by HHblits [77] as a full .a3m file (251 sequences). AlphaFold2 was run as a Google colaboratory project on GitHub, in which the MSA .a3m file and the AstaP sequence were uploaded separately (https://colab.research.google.com/github/sokrypton/ColabFold/blob/main/verbose/alphafold_noTemplates_noMD.ipynb). RoseTTAFold was run as a separate Google colaboratory project on GitHub with the same inputs as for AlphaFold2 (https://colab.research.google.com/github/sokrypton/ColabFold/blob/main/RoseTTAFold.ipynb).

### Thin-layer chromatography

Carotenoids were extracted from various holoforms of AstaP or *E. coli* membranes by the addition of a two-fold volume excess of pure acetone. Aliquots of the samples clarified by centrifugation were subjected to thin-layer chromatography on silica gel plates (Silufol, Kavalier, Czechoslovakia) using a mixture of kerosene (70% v/v) and acetone (30% v/v) for 10 min at room temperature. The results were recorded immediately after the run, to prevent oxidation of carotenoids. Rf values for different pure carotenoids were used as a reference.

### Analytical size-exclusion spectrochromatography (ASESC)

Size-exclusion chromatography with diode array detection was used to obtained spectrochromatograms of AstaP holoforms with different carotenoids, as well as to evaluate the results of carotenoid transfer between proteins and between AstaP and the membranes. To this end, samples (45 μl) were loaded on a Superdex 200 Increase 5/150 column (GE Healthcare) operated at a 0.45 ml/min flow rate using a Varian 335/Varian 363 HPLC system (Varian Inc., Melbourne, Australia). The column was pre-equilibrated with a filtered and degassed 20 mM Tris-HCl buffer, pH 7.6, containing 150 mM NaCl. During the run, absorbance in the 240-900 nm range with 1-nm steps (4 nm slit width) was recorded with a frequency of 2.5 Hz. Alternatively, ASESC was used during the final preparative SEC, in which case a Superdex 75 26/60 column (GE Healthcare) operated at a 2.6 ml/min flow rate was used. SEC profiles and absorbance spectra were extracted from the DAD data using a custom Python-based script. Spectrochromatograms were processed and visualized in OriginPro 9 (Originlab, Northampton, MA, USA). Apparent Mw values for the peaks were determined via column calibration using BSA dimer (132 kDa), BSA monomer (66 kDa), ovalbumin (43 kDa), and α-lactalbumin monomer (15 kDa).

To analyze the interaction of AstaP with liposomes, the ASESC profiles obtained from a Superdex 200 Increase 5/150 column were fractionated using a Bio-Rad 2110 fraction collector, and AstaP partition between the protein and liposome fractions was assessed by SDS-PAGE [78].

### SEC-MALS

Size-exclusion chromatography coupled with multi-angle light scattering (SEC-MALS) was done by connecting of either a Superdex 200 Increase 10/300 or a Superdex 75 10/300 (both GE Healthcare) to a UV-Vis Prostar 335 detector (Varian, Australia) and a miniDAWN detector (Wyatt Technology, USA). AstaP samples (150-300 µg added in 100-120 μl) were applied at a 0.8 ml/min flow rate to the column equilibrated with a filtered (0.1 μm) and degassed 20 mM Tris-HCl buffer, pH 7.6, containing 150 mM NaCl. Signals from the detectors were processed with ASTRA 8.0 software (Wyatt Technology, USA) using dn/dc equal to 0.185 and extinction coefficients ε^0.1%^_280nm_ of 1.04 (AstaP apoprotein), 1.44 [AstaP(CAN)] and 1.75 [AstaP(ZEA)]. The extinction coefficients for the holoforms took into account concentration of the protein determined by the Bradford assay [74]. Since these samples could be contaminated by the apoprotein, the corrected extinction coefficients could be valid only for the specific samples obtained.

### Absorbance measurements

Steady-state absorption spectra and the time-courses of absorbance changes at 550 nm were recorded as described earlier [50, 56]. For the photoconversion of the samples (actinic light for OCP^O^ → OCP^R^ photoconversion) a blue light-emitting diode (M455L3, Thorlabs, USA) with a maximum emission at 445 nm was used. The temperature of the sample was stabilized by a Peltier-controlled cuvette holder Qpod 2e (Quantum Northwest, USA). Each experiment was repeated at least three times, and the most typical results are presented.

Absorbance spectra of carotenoids in organic solvents were recorded in 1 cm quartz cuvettes on a Nanophotometer NP80 (Implen, Germany).

## Acknowledgements

The authors are grateful to Prof. Kai-Hong Zhao (Huazhong Agricultural University, Wuhan, China) for the pBAD-crtW plasmid. The study was supported by the Russian Foundation for Basic Research and the German Research Foundation joint grant (no. 20-54-12018 and no. FR1276/6-1). Size-exclusion spectrochromatograms were obtained in the framework of the Program of the Ministry of Science and Higher Education of Russia (N.N.S. and Y.B.S.). CD measurements were done at the Shared-Access Equipment Centre “Industrial Biotechnology’’ of the Federal Research Center “Fundamentals of Biotechnology” of the Russian Academy of Sciences.

## Author contributions

NNS – conceived the idea, designed the experiments and coordinated the study; YBS, NAE, NNS – expressed and purified proteins; YBS, NAE, NNS – performed experiments; YBS, EGM, TF, NNS – analyzed data and discussed the results; NNS wrote the paper with input from all authors.

## Abbreviations used

βCar: β-carotene,
ASESC: analytical size-exclusion spectrochromatography,
AstaP: astaxanthin-binding protein,
AXT: astaxanthin,
CAN: canthaxanthin,
CD: circular dichroism,
CTD: C-terminal domain,
CTDH: C-terminal domain homolog,
DA: dark-adapted,
DAD: diode array detector,
HCP: helical carotenoid protein,
ID: intrinsically disordered,
IDR: intrinsically disordered region,
IMAC: immobilized metal-affinity chromatography,
IPTG: isopropyl-β-thiogalactoside,
LA: light-adapted,
LED: light-emitting diode,
NMR: nuclear magnetic resonance,
NTD: N-terminal domain,
OCP: orange carotenoid protein,
OCP^O^: the orange dark-adapted form of OCP,
OCP^R^: the red light-adapted form of OCP,
ROS: reactive oxygen species,
SDS-PAGE: sodium dodecyl sulfate-polyacrylamide gel electrophoresis,
SEC: size-exclusion chromatography,
SEC-MALS: size-exclusion chromatography coupled to multi-angle light scattering,
ZEA: zeaxanthin.

